# Transcription termination safeguards quiescent chromatin for faithful cell-cycle re-entry

**DOI:** 10.64898/2026.03.28.714970

**Authors:** Annapoorna K. Prashanth, Anubhav Bhardwaj, Rajashekar Varma Kadumuri, Sreenivas Chavali, Venkata R. Chalamcharla

## Abstract

Quiescence is a conserved program of endurance and readiness in non-cycling cells that is fundamental to the longevity of eukaryotic lineages, from clonal microbial populations to human regenerative tissues. While this G_0_ state is universally associated with a compact chromatin organization, the active safeguards that preserve this structural template—and their necessity for future division competence and fidelity—remain unknown. Here we show, using fission yeast as a model, that chromatin structural maintenance in quiescence requires the enforcement of transcription termination by the conserved factor Ppn1^PNUTS/Ref2^. We identify a minimal disordered region in Ppn1 that resolves a physical conflict in the quiescent genome by preventing the transcriptional “eviction” of cohesin. Loss of this safeguard drives a progressive structural erosion that deregulates cyclin-dependent kinase (CDK) dynamics and causes lethal aneuploidy upon cell-cycle re-entry. Notably, this deterioration is not an inevitable terminal state; re-establishing termination via a brief Ppn1 pulse resets division fidelity by stabilizing a core *de novo* cohesin landscape. These findings redefine quiescent chromatin as an actively maintainable blueprint rather than a passive standby state established at G_0_ entry. We propose that “quiescence exhaustion” is driven by transcriptional stress eroding genomic organization, defining a structural limit to cellular longevity.

## Main Text

Quiescence is a reversible noncycling state whose conserved program of endurance and readiness supports long-term cellular survival from microbes to human stem cells (*1*). Compactly organized chromatin structure is a key feature of quiescence: Entry into quiescence is accompanied by rapid chromatin condensation and nuclear reorganization (*2, 3*), leading to widespread transcriptional silencing and tight control of cell-cycle gene expression (*4*). Despite low transcriptional activity throughout quiescence, the chromatin remains accessible to transcription factors, which modulate the expression of pro-survival genes to withstand age-induced stress, in addition to maintaining the expression of “reversibility” genes that control cellular responsiveness to physiological growth signals (*5*). Moreover, recent studies indicate that despite compaction, chromatin in G_0_ is readily primed for a robust increase in transcription and governs accurate cell-cycle re-entry when stimulated (*6, 7*).

The structural maintenance of chromosomes (SMC) complexes—condensin and cohesin—are key determinants of chromatin organization during cell division cycle and also in G_0_ (*8-11*). While recent studies reveal that quiescent cells actively replenish centromeric histones and prime promoters for reactivation (*6, 7*), the mechanisms that safeguard the global integrity of the chromatin structure during extended quiescence remain undefined. RNA polymerase II (Pol II) transcription has emerged as a central regulator of genome topology across the cell cycle, influencing the distribution and stability of SMC complexes and balancing cohesin-mediated looping with condensin-driven compaction. Interestingly, rather than transcription elongation simply displacing SMCs (*12, 13*), transcription termination factors can both oppose condensin recruitment and promote its access by clearing Pol II roadblocks (*14, 15*), while also preserving cohesin organization by preventing readthrough transcription (*16*). In fission yeast, Pol II transcription and termination factors drive heterochromatin formation (*17, 18*); this constrains chromatin interactions, shapes nuclear architecture, and is dynamically regulated to promote quiescence entry and maintenance (*19, 20*).

Cell cycle withdrawal into quiescence is governed by extrinsic signals that converge on cyclin dependent kinase (CDK) activity, and across eukaryotes, CDKs set commitment to the cell cycle and orchestrate the orderly execution of events that ensure its successful progression (*21, 22*). CDK activity, fundamentally driven by mitogenic signal-dependent transcriptional control of cyclins and their modulators, feeds back to shape the global cell-cycle transcriptional program, ensuring phase-appropriate gene expression (*23, 24*). This temporal coordination aligns major events—single-round DNA replication, mitotic entry with spindle and chromatin reorganization, and orderly mitotic exit—and safeguards fidelity at each checkpoint (*25-27*). Disrupting this timing causes arrest or lethal errors that compromise cell viability and identity. It remains unclear how quiescent cells reset fidelity in transcriptional control of the cell cycle upon reactivation from G_0_. In particular, the ability to accurately execute mitosis after prolonged G_0_ is a necessity for population expansion, but whose mechanisms are poorly understood. Using *Schizosaccharomyces pombe* (*S. pombe*)—a well-established model for both cell-cycle control and long-term quiescence (*20, 28, 29*)—we reveal that enforcing transcription termination is a fundamental requirement to preserve the G_0_ chromatin structure, which serves as a structural framework to ensure precise CDK regulation and division fidelity upon reactivation.

### Chromatin structural determinants of G_0_ division-competence

The transition from proliferation to quiescence entails extensive proteome remodeling (*4*). In *S. pombe*, nitrogen starvation induces a synchronized entry into a reversible G_0_ state, where they remain viable for weeks (*28*). To identify factors that preserve chromatin structure during this state, we isolated chromatin from G_0_ *S. pombe* cells and quantified associated proteins (the “chromatome”) relative to the whole-cell proteome using tandem mass tag (TMT) labeling and mass spectrometry. Several conserved chromatin architectural and regulatory components (*11, 14, 19, 30*)—including the cohesin and SMC5/6 complexes, DNA topoisomerase II (Top2), heterochromatin protein 1 (Swi6), and the DPS^PP1/PTW^ phosphatase complex—remained consistently enriched on quiescent chromatin (fig. S1 and Table S2).

To determine which of these candidates are required for the long-term maintenance of G_0_ chromatin organization and viability, we generated a panel of prototrophic, Tet-OFF strains to deplete representative 3xFLAG-tagged proteins specifically after the establishment of quiescence. All strains reached the 1C DNA content characteristic of G_0_ entry normally, as confirmed by flow cytometry 24 hours post-induction (fig. S2A). At this time point, anhydrotetracycline (AhT) was added to the media and maintained for three weeks to produce rapid and sustained protein depletion (fig. S2B). We monitored nuclear morphology and chromatin organization using the envelope marker Ish1–GFP and the chromatin marker H2A– mCherry in young (3 days with AhT) and aged (21 days with AhT) G_0_ cells. In wild-type controls, chromatin remained organized in its hallmark compact G_0_ structure in both young and aged cells. By contrast, depletion of the cohesin subunit Psc3, Top2, or Ppn1 (the yeast homolog of human PNUTS) compromised this stability (fig. S2C); aged cells exhibited a marked loss of chromatin sphericity—quantified by 3D shape analysis—that was not observed in young quiescent cells (fig. S2D).

Among the candidates screened, Ppn1 depletion showed the most profound effect on G_0_ maintenance, resulting in a marked loss of viability in single-cell colony-forming assays (fig. S2E). In cycling cells, Ppn1 recruits Protein Phosphatase 1 (PP1) to the nucleus for transcriptional control, mitotic progression, and genome maintenance (*31-33*). While many of these functions require the Ppn1–PP1 holoenzyme, Ppn1 also acts as a multivalent scaffold that organizes a diverse network of proteins beyond PP1 (*34, 35*). To further characterize the requirement for Ppn1 during quiescence, we generated a *ppn1*Δ null mutant and monitored its G_0_ nuclear organization at weekly intervals using the markers described above (Fig. 1A). 3D shape analysis of quiescent nuclei revealed a progressive loss of chromatin sphericity that closely paralleled the decline in cellular viability (Fig. 1, B and C). Notably, in *ppn1*Δ cells, telomeres in the characteristic Rabl configuration were already declustered by day 3 of G_0_, while centromere clustering remained largely intact (Fig. 1D). Upon reactivation, *ppn1*Δ cells showed delayed S phase entry and underwent mitosis with chromosome missegregation, cytokinesis failure, and abnormal septation—revealing broad defects in cell-cycle control (Fig. 1, C and E, and fig. S3, A to C).

**Fig. 1.**
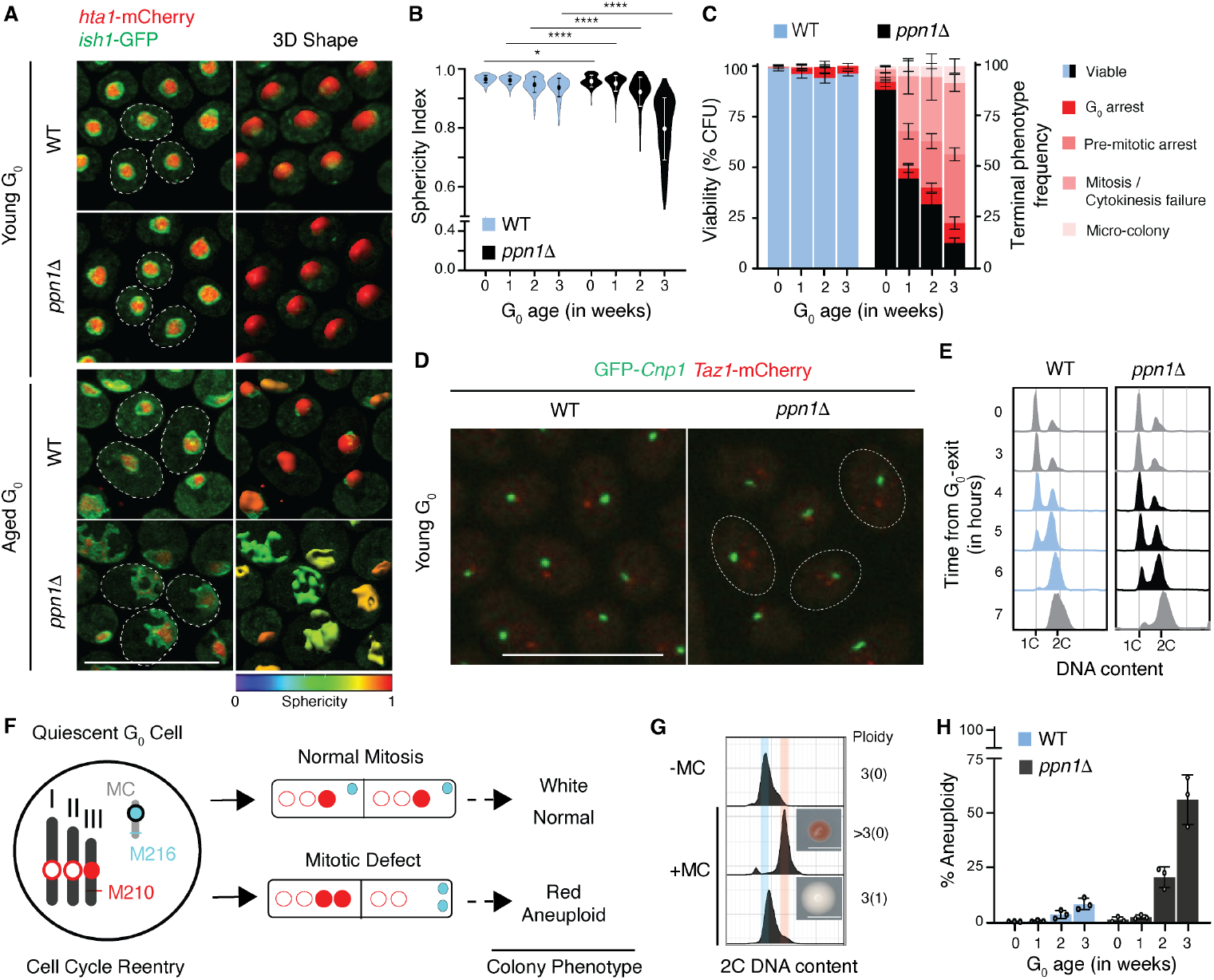
Safeguarding of quiescent chromatin structure and G_0_ viability by Ppn1. **(A)** Hta1– mCherry (chromatin) and Ish1–GFP (nuclear envelope) imaging in WT and *ppn1*Δ cells over three weeks of quiescence. Chromatin (Hta1–mCherry) and Nuclear envelope (Ish1 GFP) imaging in young (3 days after G_0_-entry) and aged (21 days after G_0_-entry) quiescent cells of wild-type (WT) and *ppn1*Δ prototrophic strains. **(B)** Chromatin sphericity loss quantified in WT and *ppn1*Δ over three weeks of quiescence (*n* > 500 cells; \*\*\*\**P* < 0.0001, \**P* < 0.05; two tailed *t*-test; three biological replicates). **(C)** G_0_ viability (CFU, colony-forming units) and terminal phenotypic analysis in WT and *ppn1*Δ over three weeks of quiescence (*n* > 500 cells; three replicates). **(D)** Telomere (Taz1–mCherry) and centromere (GFP–Cnp1) organization in young G_0_ cells; *ppn1*Δ exhibits multiple red foci, indicative of declustering, whereas WT displays a single clustered focus. **(E)** DNA content (flow-cytometry) profiles upon G_0_-exit; *ppn1*Δ displays a ∼1-hour delay in S-phase onset compared to WT. **(F)** Schematic of the Ch16 minichromosome (MC) aneuploidy assay. The MC (blue circle) carries the *ade6-M216* allele in a host background containing *ade6-M210* (red circles, endogenous chromosomes). To track segregation, one of the three regular chromosomes is depicted as a filled red circle; normal mitosis results in white colonies via interallelic complementation (+MC) whereas missegregation during the first post-quiescence division yields red colonies (–MC), primarily capturing survivable aneuploidy events where MC loss is accompanied by endogenous chromosome gain. **(G)** DNA content profiles of representative *ppn1*Δ colonies following G_0_ exit; red colonies exhibit a shift toward higher DNA content (≥3 regular:0 MC), confirming aneuploidy arising from endogenous chromosome gain rather than simple MC loss. A white colony from a ‘prototrophic’ cycling cell (lacking MC) is shown as reference. **(H)** Aneuploid survivor (red colonies) frequency with age in quiescence (*n* ≥ 3 biological replicates. Error bars indicate standard deviation (SD). Scale bars, 10 μm.

Aneuploidy, arising from segregation errors, has been observed—albeit at low frequency—in quiescent and mitotic tissue stem cells across diverse lineages (*36*). We therefore assessed aneuploidy during the first mitosis after quiescence in *S. pombe* using the well-established Ch^16^ minichromosome (MC) assay (*37*), in which retention yields white colonies and loss produces red (Materials and methods; Fig. 1F). MC loss during normal proliferation was low in both wild-type and *ppn1*Δ cells (fig. S3D). In contrast, prolonged quiescence in *ppn1*Δ cultures produced numerous small red colonies, while wild type showed only a modest age-associated increase (Fig. 1, G and H, and fig. S3E). Unexpectedly, flow cytometry revealed that these red colonies— which were expected to show a reduction in DNA content following MC loss (3:0)—actually possessed a significantly higher total DNA content than white (3:1 or 3:2) colonies (Fig. 1G). This indicates that in *ppn1*Δ cells, loss of the reporter MC is a proxy for broader aneuploidy of the endogenous chromosomes (>3:0). Because chromosome gain or loss is poorly tolerated in fission yeast (*37*), these frequencies likely underestimate the true extent of aneuploidy. Collectively, these experiments indicate that Ppn1 safeguards chromatin structural integrity during prolonged quiescence, enabling accurate first mitosis.

### Readthrough transcription-driven chromatin structural erosion in G_0_

Given Ppn1’s diverse roles as a multivalent scaffold, we performed an unbiased genetic suppressor screen to identify the specific pathway essential for quiescence. Using the sharp loss of *ppn1*Δ viability in G_0_ as a selection (Materials and methods; fig. S4A), we found that all recovered suppressors mapped to core RNA polymerase II (Pol II) or elongation factors that promote transcriptional progression *in vivo*, implicating dysregulated elongation as the driver of *ppn1*Δ inviability (Fig. 2A, and fig. S4, B and C). Consistent with this, dampening elongation using either of two well-characterized “slow” Pol II catalytic alleles or 6-azauracil (6AU) treatment—rescued *ppn1*Δ survival in quiescence (Fig. 2B). To test causality, we examined chromatin structure in these defined backgrounds; unlike *ppn1*Δ, they lacked global chromatin defects (Fig. 2C), though telomere declustering persisted in the slow-Pol II suppressor (Fig. 2D). Together, these distinctions indicate that aberrant Pol II progression is the primary driver of G_0_ inviability, whereas telomere declustering has an independent cause.

**Fig. 2.**
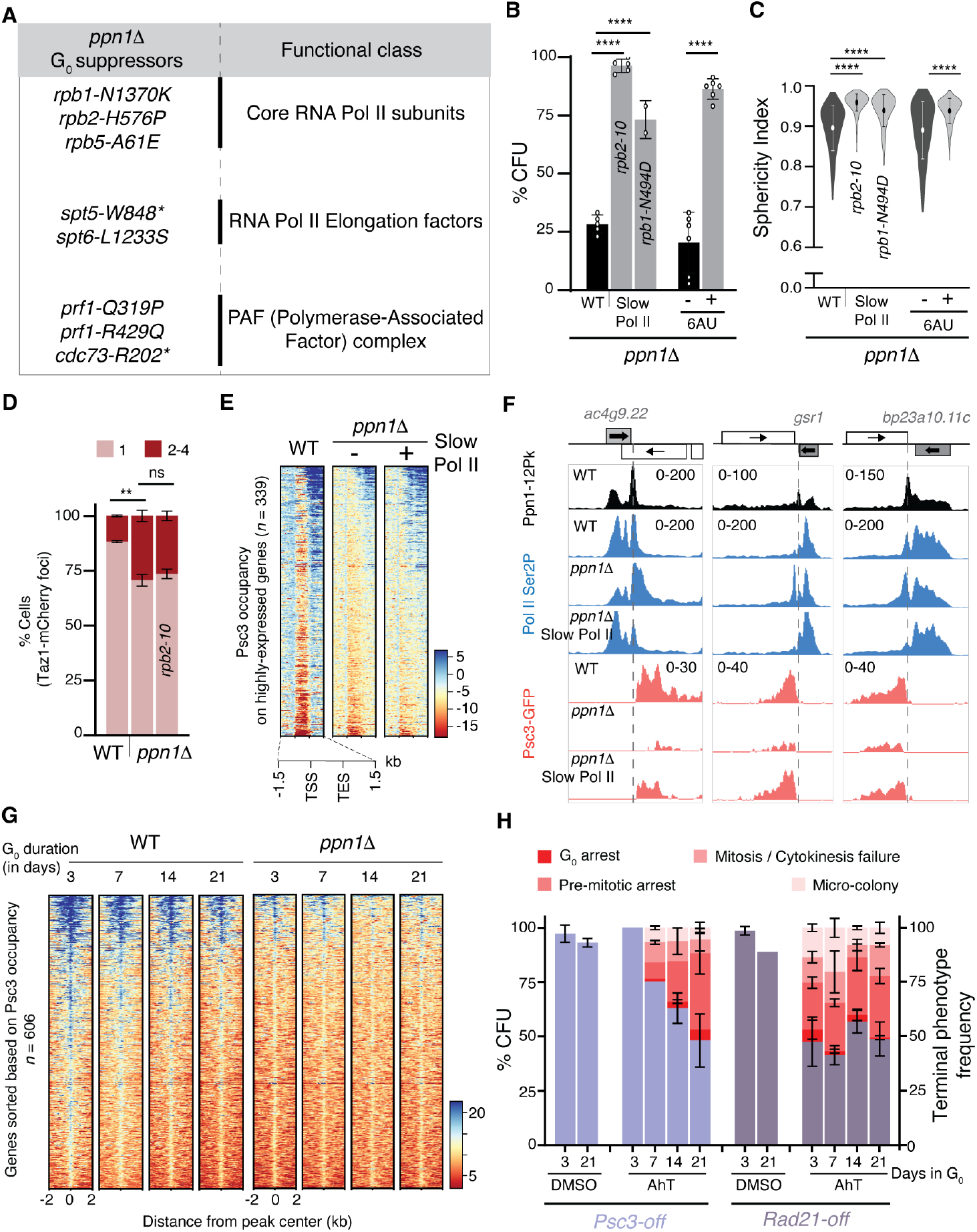
Global maintenance of G_0_ cohesin by Ppn1-dependent Pol II transcription. **(A)** *ppn1*Δ G_0_ suppressors map to core RNA Pol II subunits and positive transcription elongation regulators. **(B)** Rescue of *ppn1*Δ viability (14-day G_0_) by slow Pol II mutations or 6-azauracil (6AU) (*n* ≥ 2 biological replicates). **(C)** Suppression of chromatin distortion in *ppn1*Δ (14-day G_0_) by slow Pol II or 6AU (\*\*\*\**P* < 0.0001; two-tailed *t*-test; *n* > 350 cells; 2 biological replicates). **(D)** Quantification of Taz1–mCherry foci (3-day G_0_) showing persistent telomere declustering in *ppn1*Δ and its slow-Pol II suppressor (\*\**P* < 0.005; two-tailed *t*-test; *n* > 200 cells, 2 biological replicates). **(E)** Heatmaps of Psc3–GFP ChIP-seq at highly expressed genes (3-day G_0_). Cohesin enrichment at 3′ ends are reduced in *ppn1*Δ and restored in the suppressor (spike-in normalized). TSS, Transcription Start Site; TES, Transcription End Site. **(F)** Browser views at representative genes (3-day G_0_) show Ppn1 enrichment at the TES (gray dotted line) coincides with *ppn1*Δ readthrough and cohesin loss, both suppressed by slow Pol II. **(G)** Peak-centered heatmaps showing stable cohesin occupancy in WT over three weeks (*n* = 606 non overlapping genes), contrasted with persistent loss in *ppn1*Δ. **(H)** Conditional depletion of Psc3 or Rad21 by AhT phenocopies *ppn1*Δ viability loss and terminal morphologies (*n* = 2 biological replicates). Error bars represent SD.

To examine the transcriptional role of Ppn1 in quiescence, we performed ChIP-seq of epitope tagged Ppn1 (Ppn1-12Pk) and elongating Pol II (Ser2-phosphorylated CTD, Ser2P). Consistent with its role in elongation control and termination, Ppn1 accumulated at gene 3’ ends where Pol II typically pauses during termination; in *ppn1*Δ cells, however, the Pol II signal extended well beyond annotated termination sites (TES), indicating pervasive transcriptional readthrough (fig. S5A). Because cohesin and condensin localize near gene 3′ ends—cohesin during interphase and condensin during mitosis (*12, 13*)—we next examined how this readthrough affects their placement in quiescence. ChIP-seq of the cohesin complex using the core subunit Psc3 (Psc3 GFP) revealed that in wild-type G_0_ cells, cohesin is concentrated just downstream of the TES, predominantly at transcriptionally active loci (Fig. 2E and fig. S5B). In *ppn1*Δ, cohesin on chromosome arms was lost when elongation was unchecked, whereas dampening elongation largely preserved these 3’-end peaks (Fig. 2, E and F, and fig. S5C). Importantly, at transcriptionally repressed centromeres and telomeres, cohesin was largely unaffected by Ppn1 loss (fig. S5D), further linking the cohesin loss to aberrant transcription.

### Global G_0_ cohesin maintenance via Ppn1-dependent Pol II termination

Across prolonged quiescence, cohesin positioning on chromosome arms remained stable in wild type—reflecting the active maintenance of higher-order chromatin organization— whereas in *ppn1*Δ, the deficit at gene ends persisted without redistribution (Fig. 2G). These data indicate that the *ppn1*Δ defect likely represents a failure of cohesin retention specifically at transcribed regions. By contrast, condensin (Cut14–12Pk) in wild-type G_0_ was enriched along gene bodies upstream of gene ends (fig. S6, A to C). In *ppn1*Δ, condensin was selectively reduced on gene bodies overlapping canonical mitotic targets, while several new G_0_ sites were unaffected (fig. S6D). Notably, depleting cohesin subunits alone recapitulated *ppn1*Δ chromatin defects and broad reentry failures (Fig. 2H and fig. S2B), identifying cohesin instability as the downstream driver of the mutant phenotypes.

Our findings indicate that Ppn1 functions independently of PP1 phosphatase to maintain quiescence. Loss of Dis2, the major PP1 isoform implicated in termination (*33, 38*), only partially recapitulated *ppn1*Δ phenotypes, and mutations that abolish the PP1-bindingmotif (ABC) had no effect on viability or chromatin structure (fig. S2 and S7A; (*14*)). Systematic truncations showed that deleting the N-terminal TND (TFIIS N-terminal domain; common in elongation regulators (*14, 39*)) or the C-terminal Swd2.2-binding site that tethers Ppn1 to core cleavage and polyadenylation factors (CPF (*14, 39*)) had no effect in G_0_ (Materials and Methods; Fig. 3, A and B). By contrast, the central intrinsically disordered region (IDR), implicated in Pol II binding (*40*), proved essential: a C-terminal truncation (ΔC-long) removing a discrete IDR segment failed to localize to the nucleus in cycling cells and fully phenocopied *ppn1Δ*—loss of G_0_ viability, reduced cohesin occupancy, and pervasive Pol II readthrough—whereas a shorter truncation sparing this segment (ΔC-short) remained functional (Fig. 3, B to D, and fig. S7, B to D). Although this segment includes the PP1-binding site, PP1 binding was dispensable, implicating other essential features within the conserved IDR (fig. S7E).

**Fig. 3.**
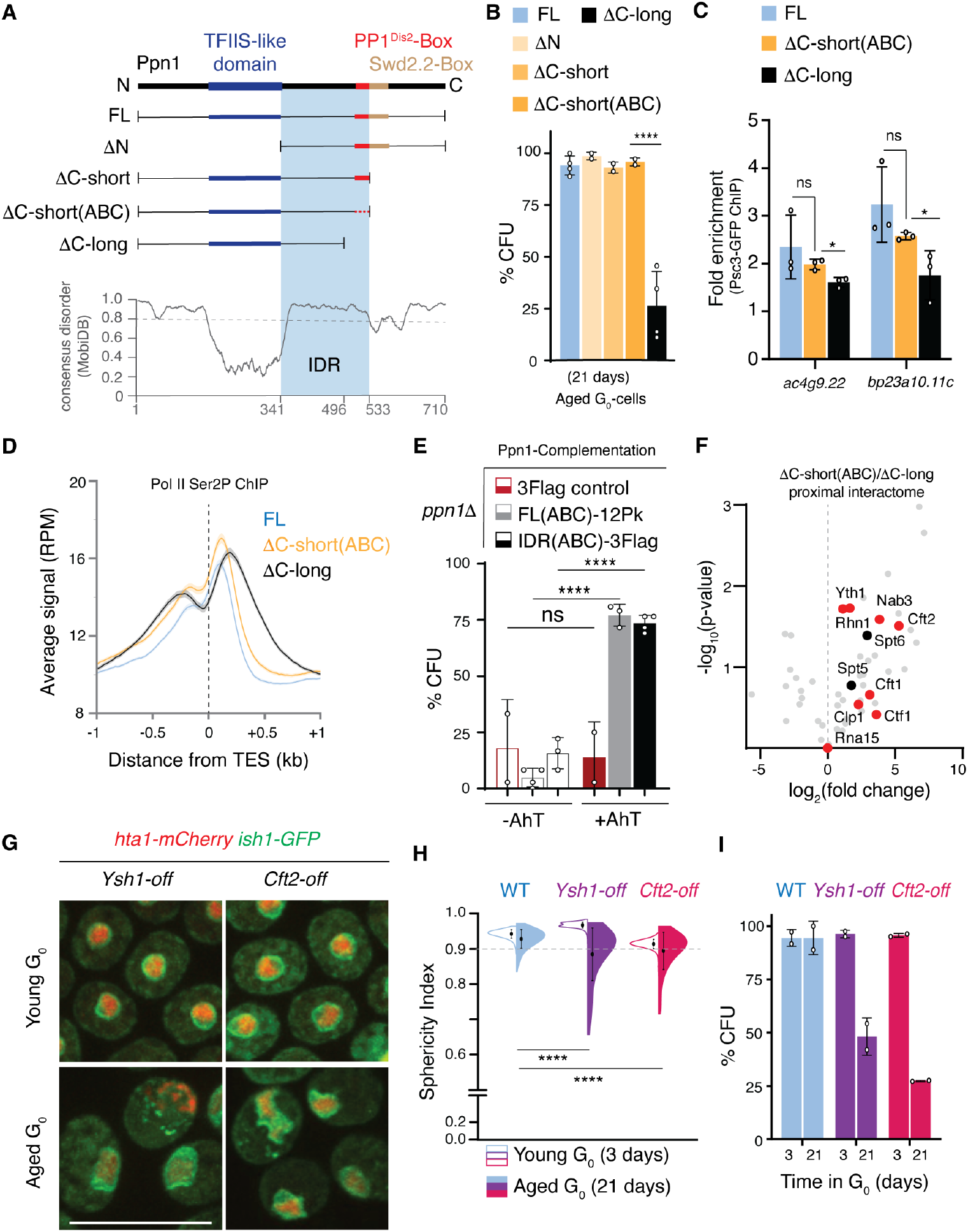
Coordination of G_0_ CPF-mediated termination by a minimal Ppn1 IDR. **(A)** Schematic of Ppn1 truncations. **(B)** G_0_ viability identifies a critical 36-amino-acid IDR segment (residues 497-532) required for survival (\*\*\*\**P* < 0.0001; two-tailed *t*-test; n ≥ 2 biological replicates). **(C)** ChIP-qPCR of Psc3-GFP (3-day G_0_) confirms cohesin loss at 3′ ends in ΔC-long (lacking IDR) but not functional ΔC-short(ABC) (*n* = 3; \**P* ≤ 0.05; two-tailed *t*-test). **(D)** Metagene ChIP-seq of elongating Pol II (Ser2P; spike-in–normalized) at 3-day G_0_. Elevated downstream occupancy in ΔC-long links the IDR to termination enforcement (*n* = 3329 genes; 2 replicates). **(E)** Inducible complementation of *ppn1*Δ G_0_ with Ppn1 full-length (FL) or IDR only (342–532 aa) variants, both carrying mutations in the PP1-binding site (ABC). Continuous induction of the IDR fully restores *ppn1*Δ viability (21-day G_0_) (\*\*\*\**P* < 0.0001; ns, not significant; two-tailed *t*-test; *n* ≥ 2 biological replicates). **(F)** TurboID-based proximity labeling comparing functional (ΔC-short(ABC)) and non-functional (ΔC-long) variants identifies IDR-dependent interactors, including Cleavage and Polyadenylation Factors (CPF; red) and Pol II-binding proteins (black). **(G-I)**, Targeted depletion of CPF subunits Ysh1 or Cft2 phenocopies *ppn1*Δ chromatin and survival defects. Scale bars, 10 μm; \*\*\*\**P* < 0.0001; two-tailed *t*-test; error bars, SD.

### Coordination of CPF-mediated termination by a minimal Ppn1-IDR

To test sufficiency, we ectopically expressed either full-length Ppn1 or the Ppn1-IDR (aa 342 532) in *ppn1Δ* cells. Remarkably, the IDR alone—with or without the PP1-binding site—was sufficient to restore full G_0_ viability, mirroring the rescue observed with the full-length protein (Fig. 3E and fig. S8A). To define the chromatin landscape supporting this recovery, we performed ChIP-seq of Psc3–GFP upon full-length Ppn1 complementation. This identified 127 significantly and reproducibly enriched cohesin peaks (fig. S8B and Table S3). While this represents a subset of the 501 native peaks found in wild-type G_0_ cells, this core landscape was sufficient for the full recovery of viability. Spatial concordance analysis revealed that 93% of these core peaks (117/127) matched the exact target gene or were located within 5 kb of the nearest native target locus (fig. S8, C and D). The functional importance of this architectural core was further validated by IDR-only complementation, which showed high concordance with the core set (51% genomic overlap; fig. S8B). Notably, a small number of peaks found upon complementation were absent from both wild-type and *ppn1Δ* cells, indicating that they arise *de novo* at transcribed loci upon the restoration of termination (fig. S8, E and F). These patterns demonstrate that G_0_ structure is maintained by an active balance between continuous cohesin turnover and Ppn1-dependent stabilization. While the cohesin loading machinery remains functional during quiescence consistent with the observation that acute Mis4 inhibition disrupts nuclear separation at exit (*41*)—Ppn1-dependent termination serves as the essential safeguard that prevents transcriptional eviction from overwhelming the core landscape.

To define the IDR contacts that mediate termination during quiescence, we mapped IDR dependent partners by TurboID proximity labeling in G_0_ cells. Mass spectrometry showed that ΔC-short(ABC) lost multiple CPF contacts—including Dis2 and Swd2.2—consistent with disruption of the CPF–phosphatase interface, yet retained a subset of CPF subunits (Fig. 3F, fig. S9, and Table S4). ΔC-long further eroded these interactions, with a complete loss of Cft2 (CPF nuclease module) and markedly reduced association with elongation factors Spt5 and Spt6. These results indicate that residues 497–532 within the IDR are critical for CPF–Pol II coordination during termination (Fig. 3F). Depletion of Cft2 or the endonuclease Ysh1 in quiescent cells phenocopied Ppn1 loss—showing progressive chromatin deterioration and G_0_ inviability underscoring that canonical polyadenylation site (PAS)-dependent termination is required to maintain the quiescent state (Fig. 3, G to I). Together, these findings define a minimal IDR module that enforces Pol II termination and sustains cohesin-dependent chromatin stability in G_0_, independently of PP1.

### Cohesin restoration and structural repair in aged G_0_ cells

Next, we investigated whether the architectural collapse of the G_0_ nucleus is reversible through the timed restoration of Ppn1. Using an AhT-inducible Tet-ON system, we induced full-length Ppn1 either for a brief 4-hour window in G_0_ or just before reactivation (Fig. 4A). Remarkably, 4 hours of induction in G_0_ was sufficient to restore full viability even in deep quiescent cells, whereas induction after exit failed to rescue despite robust expression prior to S-phase (Fig. 4B and fig. S10A). Only complementation in G_0_ itself prevented the abnormal division phenotypes typically seen during reactivation (fig. S10B).

**Fig. 4.**
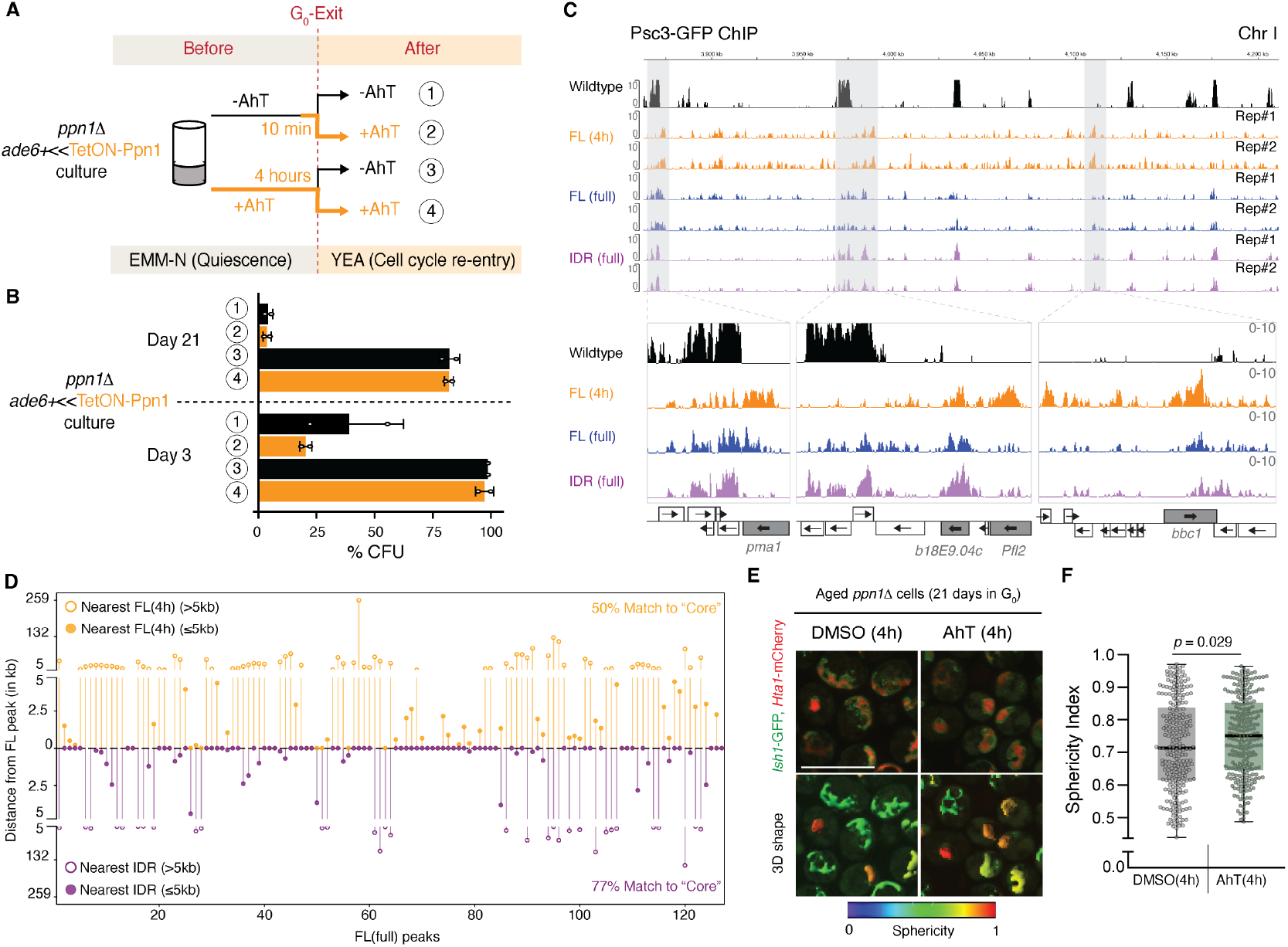
Restoration of a “core” G_0_ cohesin landscape and a division-blueprint. **(A)** Schematic of AhT-inducible complementation assay. **(B)** Rescue of *ppn1*Δ viability requires Ppn1 expression specifically during quiescence; post-exit induction is insufficient (*n* = 2 biological replicates). **(C)** Restored Psc3–GFP ChIP-seq (3-day G_0_) profiles following complementation with full-length (FL) or IDR-only Ppn1. Browser views compare continuous (3-days) induction with an acute (4 hours) pulse administered prior to harvest; profiles are spike-in–normalized and background-subtracted. **(D)** Spatial concordance of the cohesin template required for quiescence. Proximity analysis (3-day G_0_) of IDR (full induction) and FL (4 h pulse) datasets relative to the 127 core reference loci (see fig. S8, B and D). High-frequency matching (77% for IDR; 50% for 4 h) within 5 kb indicates that this shared core, plus condition-specific sites (see fig. S8, E and F), is sufficient for structural integrity and survival. **(E)** Rapid architectural restoration in aged (Day 21) G_0_ cells. Hta1–mCherry (chromatin) and Ish1–GFP (nuclear envelope) imaging reveals that 4-h Ppn1 induction within the quiescent state reverses age-associated nuclear deformation. **(F)** Chromatin sphericity quantification. 3D shape analysis from Hta1–mCherry signal in aged G_0_ cells (*n* > 200 cells) reveal that Ppn1 restoration rescues chromatin deformation and improves compaction compared to DMSO control. Scale bars, 10 µm.

To determine if this rescue reflects a rapid “repair” of the cohesin landscape, we performed Psc3 GFP ChIP–seq following the 4-hour G_0_ induction. We identified 202 significant and reproducible peaks; while exact coordinate overlap with the core functional set was limited (16.8% exact match; fig. S8B), representative genome browser tracks revealed a low, broadly distributed cohesin signal that likely represents an “in-progress” re-establishment of the architectural framework (Fig. 4C). Spatial analysis revealed that 51% of the re-established peaks were within 5 kb of the “core” target set during this brief repair window (Fig. 4D and fig. S8, D to F). This rapid recovery coincides with a marked improvement in chromatin structure (Fig. 4, E and F), indicating that G_0_ maintenance is an active, ongoing process where basal transcription continuously directs the building of the cohesin landscape. In the absence of Ppn1, this same basal activity becomes destructive, necessitating a Ppn1-dependent “repair” window to reestablish a functional chromatin state before the cell attempts to divide.

### Uncoupling of re-entry commitment from cell-cycle progression

Given the severe structural deterioration in *ppn1*Δ, we asked if the failure to re-enter the cell cycle stems from a collapse of the reactivation transcriptional program. One hour after release, *ppn1*Δ failed to properly induce a significant fraction of genes that are dynamically regulated in wild-type cells during re-entry. These included the re-entry essential factor *sam1* (S-Adenosylmethionine Synthetase (*42*)) and genes required for centromere reassembly (Fig. 5A and fig. S11, A to C). Besides, in both genotypes, most canonical G_1_ genes characteristic of cycling cells remained unresponsive, indicating that re-entry uses a distinct transcriptional program (fig. S12). By 4 hours post-release, the defects in *ppn1*Δ expanded to include DNA-replication licensing factors (Fig. 5B and fig. S11D), consistent with the observed delay in S-phase onset. Notably, the transcriptomes of *ppn1*Δ and cohesin-depleted cells showed striking similarity upon reactivation (fig. S11, E and F), indicating that the structural failures established in G_0_ dictate a shared failure of the reactivation program.

**Fig. 5.**
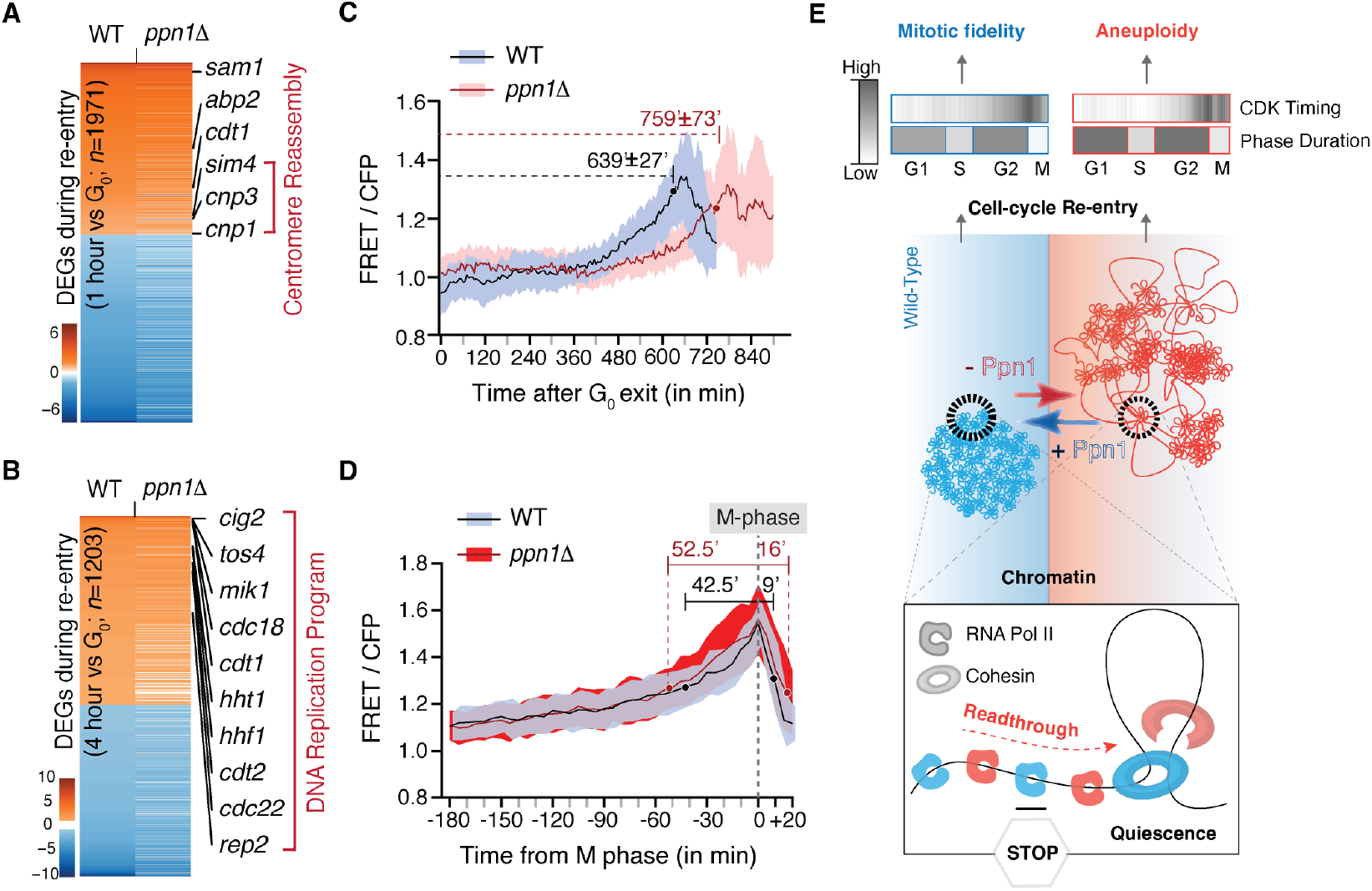
Ppn1-controlled reactivation template for faithful first mitosis. **(A, B)** Transcriptomic heatmaps of differentially expressed genes (DEGs) during exit (3-day G_0_). *ppn1*Δ displays widespread misexpression of pre-replicative and early S-phase genes. **(C, D)** Time course of CDK activity measured by Eevee-spCDK. Mean of FRET/CFP ratio of Eevee-spCDK in cells released from 3-day G_0_ reveal uncoupled CDK activation/deactivation dynamics in *ppn1*Δ. While WT cells exhibit sharp, coordinated oscillations, *ppn1*Δ cells show prolonged and erratic CDK activity primarily during G_2_ (**C**) and M (**D**). Error bars, SD. **(E)** Model comparing WT (blue) and *ppn1*Δ (red) cells. In WT, the Ppn1 safeguard preserves cohesin and chromatin compaction by enforcing transcription termination, thereby maintaining a poised state. Reactivation triggers coordinated CDK dynamics and faithful mitosis. In *ppn1*Δ, transcriptional readthrough evicts cohesin, eroding chromatin organization and uncoupling CDK control, resulting in catastrophic mitotic errors and aneuploidy. Reversible arrow denotes chromatin structural decay and restoration via Ppn1 re-expression in quiescence.

To assess the functional consequences of this collapse, we monitored cyclin-dependent kinase (CDK) activity using a FRET biosensor (*43*) (Materials and Methods; Fig. 5, C and D). Wild-type CDK dynamics were remarkably consistent between proliferating and re-entering populations, both featuring a characteristic biphasic rise through G_2_, a sharp mitotic peak, and rapid inactivation. In contrast, *ppn1*Δ cells exhibited a significant delay in the ramp from G_0_ exit to the G2–M inflection also called the ‘change point’ (759 ± 73 min vs. 639 ± 27 min in wild type). Furthermore, the ascent to the mitotic peak was steeper and the high-CDK dwell was nearly doubled (∼16 min vs. ∼9.6 min), indicating a profound loss of robustness in the cell-cycle control network. Single-cell traces were markedly heterogeneous (fig. S13), with ∼50% exhibiting distorted trajectories that correlate with the reduced colony-forming capacity of the mutant. This compromise at mitotic entry and exit results in uncoordinated mitotic events, including cytokinesis and septation (*44, 45*), providing a unifying explanation for the terminal morphologies, and aneuploidy observed in *ppn1*Δ survivors. Taken together, these findings show that chromatin structure in prolonged G_0_ is maintained by Ppn1-dependent cohesin retention at gene ends, and that the active preservation of this framework is a strict prerequisite for faithful mitotic progression upon exit.

## Discussion

Our study identifies a genome-scale safeguard that preserves the functional organization of chromatin through prolonged quiescence (Fig. 5E). We demonstrate that Ppn1-dependent transcription termination maintains cohesin positioning, sustaining a structural framework that supports gene regulation and ensures the fidelity of the first mitosis. While prior work emphasized locus-specific priming by factors such as RSC and Mis18 (*6, 7*), we introduce a paradigm in which division competence and fidelity are preserved via the active maintenance of SMC complexes. Unlike condensin, the principal driver of G_0_ compaction, cohesin’s role in quiescence has remained enigmatic. While it appears sparse in budding yeast (*3, 9*), cohesin is tightly regulated in mammalian stem cells to enforce quiescence-specific programs (*10, 46, 47*). In fission yeast, a distinct principle emerges: the quiescent genome is not a static entity, but a dynamically maintained state requiring constant termination to counteract transcriptional displacement of cohesin. This safeguard ensures that the “arrested” state is not a mere metabolic standby, but a period of continuous structural preservation necessary for robust CDK control and the prevention of age-dependent aneuploidy.

Mechanistically, termination in quiescence is enforced by a minimal Ppn1 IDR that functions independently of the PP1 phosphatase. In cycling cells, PP1 is the central effector of termination, acting via Spt5 dephosphorylation to slow elongation or promote Pol II dimerization (*33, 38, 40*). In G_0_, however, PP1 contacts proved dispensable: a discrete IDR segment was necessary and sufficient to coordinate CPF–Pol II and stabilize the core cohesin landscape. Our findings reveal that even the low basal transcriptional activity of G_0_ is sufficient to drive structural decay if this safeguard is absent. Such vulnerability to transcriptional readthrough has been observed during viral infection and cellular stress (*16, 48*), and may be especially consequential in quiescent stem cells (*49*). In these long-lived cellular lineages, impaired termination could compromise reversibility, promote senescence, or predispose the genome to instability by allowing cumulative transcriptional interference to erode the structural foundation required for future divisions.

Classical models separate cell cycle commitment from progression (*50*). In mammalian systems, an E2F-driven switch is widely viewed as irreversibly licensing smooth progression (*51, 52*). Here, however, *ppn1* and cohesin mutants achieved commitment yet failed progression, as distorted CDK trajectories led to mitotic failure. The pronounced heterogeneity of single-cell CDK traces signals a fundamental loss of robustness in the control network (*53*). Crucially, our finding that structural restoration must occur within the G_0_ state—rather than during reactivation—redefines the nature of division competence. Cell-cycle commitment is not a guarantee of success; faithful progression is strictly contingent upon a functional chromatin organization established and actively protected during arrest. Given the conservation of the Ppn1 Pol II-CPF axis, this active safeguard likely represents a broadly shared principle used by quiescent cells across evolution to ensure readiness for an uncertain future.

## Acknowledgments

We gratefully acknowledge Shiv Grewal (NCI/NIH, USA), Jürg Bähler (University College London, UK), François Bachand (Université de Sherbrooke, Canada), Shigeaki Saitoh (Kurume University, Japan), Vincent Vanoosthuyse (ENS de Lyon, France) and Purnima Bhargava (CSIR-CCMB, India) for generously providing strains. We are indebted to Hironori Sugiyama (Institute of Science, Tokyo, Japan) for expert guidance with FRET image acquisition and to Jyotsna Dhawan and Imran Siddiqi (CSIR-CCMB, India) for thoughtful discussions. We thank the CSIR-CCMB Microscopy, Sequencing, Proteomics, and Flow Cytometry Facilities and their staff for technical support. We also thank Ramakanth Chirravuri-Venkata (CSIR-CCMB, India; laboratories of Tej Sowpati and Karthik Tallapaka) for RNA-seq data analysis, Tulasi Nagabandi (CSIR-CCMB, India) for next-generation sequencing library construction, and Monica Raman and Aarthi Sukumar (VRC lab) for experimental assistance.

## Funding

Council of Scientific and Industrial Research FIRST grant MLP-0160 (VRC)

Council of Scientific and Industrial Research FBR grant MLP-0149 (VRC)

Council of Scientific and Industrial Research Fellowship (AKP)

Core funding from CSIR-Centre for Cellular and Molecular Biology (AKP and VRC)

Core funding from Indian Institute of Science Education and Research Tirupati (SC and RVK)

Ramalingaswami Re-entry Fellowship (BT/RLF/Re-entry/05/2018) from Department of Biotechnology, Government of India (SC)

Core Research Grant (CRG/2023/004691) from Anusandhan National Research Foundation, Government of India (SC)

## Author contributions

Conceptualization: AKP, VRC

Methodology and Investigation: AKP, AB

Formal analysis and Visualization: AKP, VRC

ChIP-seq computational analysis – raw data processing, and generation of metagene profiles and heatmaps: RVK and SC

Supervision and Funding acquisition: VRC, SC

Data curation: AKP

Project administration: VRC

Writing – original draft: AKP, VRC

Writing – review & editing: AKP, AB, RVK, SC, VRC

## Competing interests

Authors declare that they have no competing interests.

## Data and materials availability

All data supporting the findings of this study are available within the paper and/or the Supplementary Materials. Raw next-generation sequencing data (ChIP-seq, RNA-seq and whole-genome seq) have been deposited in the European Nucleotide Archive (ENA) under accession PRJEB101458. Microscopy datasets have been deposited in the Systems Science of Biological Dynamics (SSBD) repository under DOI [https://doi.org/10.24631/ssbd.repos.2025.11.483]. Reagents (strains and plasmids) are available from the lead contact upon request.

## Notes

https://www.ebi.ac.uk/ena/browser/text-search?query=PRJEB101458

### Competing Interest Statement

The authors have declared no competing interest.

