## Supplementary material for "Transcription termination safeguards quiescent chromatin for faithful cell-cycle re-entry": Supplemental File.pdf

Annapoorna K. Prashanth *et al.*

**This PDF file includes:**

Materials and Methods
Figs. S1 to S13
Legends for Tables S1 to S5
References (#54 - 82)

**Other Supplementary Materials for this manuscript include the following:**

Table S1 (Microsoft Excel format).
Table S2 (Microsoft Excel format).
Table S3 (Microsoft Excel format).
Table S4 (Microsoft Excel format).
Table S5 (Microsoft Excel format).

### Materials and Methods

#### Strain construction

The prototrophic wild-type *Schizosaccharomyces pombe* 972 $h^-$ strain (JB972; gift of J. Bähler) served as the parental background for all strains used in this study. Gene disruptions and epitope tagging were carried out using PCR-based methods (*54, 55*), using pFA6a plasmids as DNA templates (gifts of S. Grewal; originally described in (*54*)), or by standard genetic crosses with heterothallic laboratory strains. A complete list of strains and oligonucleotides is provided in Table S1. Slow RNA Pol II mutants *rpb1* N494D (FBY2324; gift of F. Bachand (*56*)) and *rpb2*-P100S (SPVC641; gift of S. Grewal (*57*)), as well as the strain carrying *ppn1* mutations in the PP1 phosphatase binding site (yVV1997) (gift of V. Vanoosthuyse), were crossed into prototrophic, isogenic backgrounds.

Tet-Off strains were generated by integration of pDM291-tetR′-tup11Δ70 (gift from Michael Nick Boddy (Addgene plasmid # 41028; http://n2t.net/addgene:41028; RRID:Addgene_41028), linearized with AfeI, at the *ura4* locus of JB972 to produce SPC850. Fluorescent markers were introduced by sequential C-terminal tagging: first, *Ish1* was tagged with GFP using a GFP–KanMX cassette PCR-amplified from a plasmid gifted by S. Grewal (originally described in (*54*)); KanMX was subsequently swapped to NatMX, and *Hta1* (histone H2A) was then tagged with mCherry using an mCherry–HygMX cassette PCR-amplified from a plasmid from Addgene (gift from Jian-Qiu Wu (Addgene plasmid # 105156; http://n2t.net/addgene:105156; RRID:Addgene_105156). Transformants were selected sequentially on G418 and Hygromycin, and finally on nourseothricin after the marker swap, to generate SPC1087. For most Tet-Off alleles, the endogenous promoter was replaced by homologous recombination with a PCR-amplified *tetO-Pcyc1–3×FLAG* cassette (a gift from Michael Nick Boddy (Addgene plasmid # 52689; http://n2t.net/addgene:52689; RRID:Addgene_52689)). For *ppn1*, a PCR- amplified *tetO-Pcyc1* cassette (gift from Michael Nick Boddy (Addgene plasmid # 41023; http://n2t.net/addgene:41023; RRID:Addgene_41023)) was transformed into a strain carrying *ppn1* tagged at the C terminus with *12Pk* epitope (amplified from Addgene # 49177; gift from Eishi Noguchi; http://n2t.net/addgene:49177; RRID:Addgene_49177)), and the resulting strain was crossed with SPC1087 to yield the Tet-Off *ppn1* strain.

Tet-On constructs for *ppn1* complementation were generated using modified stable integration vectors. pAV0661 (gift from Sophie Martin; Addgene plasmid #133476; http://n2t.net/addgene:133476; RRID:Addgene_133476) was first altered to create pSPC49, in which *act1–sfGFP* was replaced with *P3nmt1.cyc1T*. This backbone was then modified to generate pSPC71 (with *tetO-Pcyc1*) and pSPC72 (with *tetO-Pcyc1–3×FLAG-Tadh1*). Full-length *Ppn1*–12Pk (WT or ΔPP1-ABC mutant) was amplified from cDNA synthesized from total RNA (extracted with the NucleoSpin RNA kit, Macherey-Nagel #740955) using a gene-specific primer and PrimeScript reverse transcriptase (Takara #6110), and cloned into pSPC71 to yield pSPC76 and pSPC81 respectively. pSPC72 was used as the backbone for cloning C-terminal 3×FLAG-tagged IDR fragments (WT or ΔPP1-ABC mutant), resulting in pSPC77 and pSPC78 respectively. All Tet-On plasmids were linearized with RsrII and integrated at the *ade6* locus of SPC984 (a derivative of the ade6-D19 strain; NBRP #FY38502), which itself carries pDM291-tetR′-tup11Δ70 (gift from Michael Nick Boddy (Addgene plasmid # 41027; http://n2t.net/addgene:41027; RRID:Addgene_41027)) at *ura4*. Each construct was crossed with a *ppn1Δ*::NatMX6 strain to generate complementation strains (SPC987, SPC988, SPC1114, SPC1137, SPC1161, and SPC1162), which were freshly isolated for all assays.

Additional strains were constructed for specific assays. For aneuploidy experiments, a prototrophic $Ch^{16}$-bearing strain (SPC369) was generated by crossing SPDF1652 (gift of S. Grewal) with an *ade6-210* strain of opposite mating type. A mutant derivative lacking *ppn1* (SPC455) was created from SPC369 by PCR-based gene disruption and selection for $Ch^{16}$-based ade6 complementation. To generate the NLS–GFP control used to assess nonspecific background in GFP ChIP–seq, the NLS–GFP (GFP with a nuclear localization signal) cassette together with its upstream constitutive *tdh1* promoter was excised from pAV0478 (gift from Sophie Martin; Addgene plasmid #133498; http://n2t.net/addgene:133498; RRID:Addgene_133498) was introduced into pAV0752 (gift from Sophie Martin; Addgene plasmid #133486; http://n2t.net/addgene:133486; RRID:Addgene_133486) by SmaI–KpnI digestion to create pSPC67; pSPC67 was then linearized with AfeI and transformed into wild-type prototroph JB972, yielding the integrant SPC782 (NLS–GFP). For proximity-dependent biotinylation, the TurboID–3×Myc–kanMX6 cassette (gift from François Bachand (Addgene plasmid # 126049; http://n2t.net/addgene:126049; RRID:Addgene_126049)) was PCR-amplified and integrated at the C terminus of full-length or truncated *ppn1* alleles in JB972; truncations were generated by SOEing PCR to fuse N-terminal fragments with either TurboID-tagged or 3Flag-tagged C-terminal halves; ΔN (342–710 aa) removes the N-terminal TND motif; ΔC-short (1–532 aa) removes the C-terminal Swd2.2-binding region; ΔC-short(ABC) additionally disrupts PP1 phosphatase-binding motifs (V508A, W510A, V519A); ΔC-long (1–496 aa) deletes an intrinsically disordered region (IDR) segment encompassing the PP1-box. For CDK activity monitoring, the phototrophic  Eevee-spCDK FRET biosensor strain HS048 (NBRP #FY48348 (*43*)) was crossed with a *ppn1Δ*::NatMX6 strain to generate isogenic wild-type and mutant biosensor strains, which were used to monitor CDK FRET activity upon quiescence–exit.

**$G_0$ induction, survival, and perturbation assays**

Quiescence was induced as described previously (*28*). All strains were prototrophic; for each experiment involving *ppn1*Δ mutants or truncation alleles, strains were freshly generated by crossing to wild-type prototroph. Cultures were shifted from EMM to nitrogen-free medium (EMM–N) at 1–2 × $10^6$ cells/ml and  maintained at 30 °C. At defined time points from Day 0 (24 hours after the switch) to 3 weeks, as required for each experiment, 10 µl of culture was spotted onto YEA plates. Single cells from the inoculum center were isolated by micromanipulation (Singer MSM400), and viability was scored by colony-forming ability, as described (*20*). Quiescent cells were also released into YEA (rich medium) to monitor cell-cycle re-entry and regrowth.

For Tet-controlled assays, anhydrotetracycline hydrochloride (AhT, 2.5 µg/ml; Abcam ab145350) was added 24 hours after nitrogen starvation, after quiescence had been established. Media were replenished with AhT every 3 days, and treated cultures were  analyzed by viability assays, flow cytometry, and western  blotting. Complementation experiments followed the same regimen: AhT was supplied either throughout quiescence (Full, in EMM-N liquid medium) or briefly before $G_0$ exit (for 4 hours or for 10 min before the time of release into YEA). Viability was scored on YEA plates with or without AhT, and terminal morphologies were examined in both plate and liquid cultures (see *Microscopy*). For chemical suppression, quiescent cultures were treated with 40 µM 6-azauracil (6AU; Sigma A1757) added to EMM–N 24 hour after induction and replenished every 3 days.

**Flow cytometry**
DNA content was measured by flow cytometry as described (*58*). Cells (~1 OD) werefixed in 70% ethanol, sonicated at 21% amplitude for 10s, treated with 0.1 mg/ml RNase A (Invitrogen 12091021) in 50 mM sodium citrate (pH 7.0), stained with 2 µM Sytox Green (Invitrogen S7020), and analyzed using an LSRFortessa (BD Biosciences). Data were processed with FlowJo_v10.10.0.

For aneuploidy analysis, red and white colonies from the $Ch^{16}$ assay (rationale below) were collected directly from YE plates and processed in parallel with controls. Wild-type prototrophic colonies lacking $Ch^{16}$ served as the euploid reference (3:0; three endogenous chromosomes only). White colonies from $Ch^{16}$ strains, which retained the minichromosome through normal segregation, corresponded to 3:1 (three endogenous plus $Ch^{16}$). Red colonies represented cells that lost $Ch^{16}$ during the first mitotic division but gained an extra endogenous chromosome, yielding >3:0. These colonies consistently showed higher DNA content than both euploid (3:0) and $Ch^{16}$-retaining (3:1) controls. Because red colonies reverted rapidly to euploidy in liquid YEA, DNA content was measured directly from freshly isolated colonies; small red colonies were pooled to obtain sufficient material.

**Quantitative chromatome profiling**
Chromatin enrichment was performed as described by (*59*) with modifications. Three independent cultures of the wild-type prototrophic strain JB972 were used, including one expressing nuclear GFP to monitor enrichment specificity. Cells (~500 OD) were harvested after 2 days in quiescence (see *$G_0$ induction, survival, and perturbation assays*), fixed with 3% paraformaldehyde (30 min, room temperature), washed with PBS, and crosslinked with 10 mM DMA (45 min, room temperature). Pellets were incubated sequentially in Buffer B (100 mM PIPES-KOH pH 7.4, 0.1 M EDTA-KOH pH 8.0, 0.1% $NaN_3$, 10 mM DTT; 20 min) and Buffer C (50 mM potassium phosphate pH 7.0, 1.2 M sorbitol, 1 mM β-mercaptoethanol, 0.5 mM $MgCl_2$; 20 min). Spheroplasting was carried out in Buffer C with an enzyme mixture containing zymolyase-100T (1 mg/ml; Nacalai Tesque 07665-55), β-glucuronidase (1000 U; Sigma-Aldrich SRE0095), and chitinase (3U; Sigma-Aldrich C8241) for 45 min at 32 °C with gentle shaking until 80–90% conversion. Spheroplasts were collected, washed, and lysed in Buffer E (300 mM KCl, 50 mM HEPES-KOH pH 7.5, 2.5 mM $MgCl_2$, 0.1 mM $ZnSO_4$, 2 mM NaF, 0.5 mM spermidine, 10 mM glycerol-2-phosphate, 0.1 mM sodium orthovanadate, EDTA-free protease inhibitors) with 0.5% Triton X-100 at 4 °C until ~90% lysis. Chromatin-enriched pellets were recovered by sucrose cushion centrifugation (30% sucrose, SW Ti41 rotor, 11,100 rpm, 15 min, 4 °C), treated with TEE buffer (10 mM Tris-HCl pH 8.0, 1 mM EDTA, 0.5 mM EGTA, protease inhibitors) and RNase A (30 min, 37 °C), and subjected to alternating SDS–urea washes (2% SDS/TEE and 7.5 M urea/TEE) before final resuspension in TEE. Enrichment efficiency was validated by western blotting for depletion of GFP (Invitrogen A11122) and retention of histone H3 (Abcam ab1791).

For total proteome analysis, 50 OD from the same quiescent cultures (one JB972 with NLS-GFP and two without) were processed using the iST-NHS kit (PreOmics P.O.00026) following the manufacturer's protocol. Cells were lysed with glass beads, proteins were quantified by Qubit assay (Invitrogen), and 75 µg of lysate was used for digestion. Chromatome-enriched pellets were processed in parallel: 150 µg protein was solubilized in PreOmics LYSE-NHS buffer, reverse-crosslinked at 95 °C, treated with Benzonase (2500 U; Santa Cruz Biotechnology sc-202391), and sonicated (Covaris S220). Seventy-five micrograms per sample was then processed with the PreOmics iST-NHS workflow. Digestion and labeling were performed with a TMTsixplex Isobaric Mass Tagging kit (Thermo Fisher Scientific 90064), with three channels allocated to total proteome and three to chromatome samples. Pooled TMT-labeled peptides (100 µg) were fractionated by high-pH reversed-phase HPLC (Agilent Extend C18 column) using 20 mM ammonium formate (pH 10) in water (solvent A) and 20 mM ammonium formate (pH 10) in acetonitrile (solvent B). The 135-min gradient was as follows: 0–20 min, 1–15% B; 20–100 min, 15–60% B; 100–110 min, 60–80% B; 110–120 min, 80–100% B; 120–125 min, 100–1% B; 125–135 min, 1% B. Fractions were collected between 3 and 125 min and concatenated into 12 pools for LC–MS/MS analysis.

### Protein Extraction and Western Blotting

Yeast cells (~ 10 OD) expressing epitope tagged proteins were lysed with glass beads using a bead beater (Unigenetics), and protein was recovered by standard TCA precipitation and dissolved into SDS sample buffer prior to resolution in a polyacrylamide gel. Primary antibodies anti-FLAG M2 (Sigma F1804) and anti-V5 (Bio-Rad MA1360) were used for western blotting analyses. Secondary detection used HRP-conjugated antibody anti-mouse (Sigma A4416).

### Microscopy

#### *Chromatin shape analysis*

Live-cell imaging was performed on approximately $1 \times 10^7$ cells (0.5 mL culture),pelleted and resuspended in 20 µL EMM–N. A 1–2 µL aliquot was spotted onto an agar pad of the same medium under a coverslip, and imaging was completed within 15 min of sample preparation. Confocal z-stacks were acquired at 0.1 µm intervals using a 100× oil-immersion objective on an Olympus FLUOVIEW FV3000 by FV31S-SW software. Images were processed in Fiji to generate maximum-intensity projections (*60*), and 3D chromatin surfaces were reconstructed from the Hta1–mCherry signal using the Surfaces module in Imaris v8.4.0 (Bitplane), from which sphericity scores were calculated. The nuclear envelope and chromatin were visualized with Ish1–GFP (488 nm excitation, 507 nm emission) and Hta1–mCherry (594 nm excitation, 610 nm emission), respectively.

#### *Terminal morphology analysis*

Terminal morphology was assessed in parallel with viability assays in both plate and liquid media. On YEA plates, cells that failed to generate visible colonies were revisited and imaged using a CMOS camera mounted on the Singer MSM400 micromanipulator. Inviable cells were categorized into characteristic terminal phenotypes ($G_0$ arrest, pre- mitotic arrest, mitosis/cytokinesis failure, and microcolonies). Fractions of each category were quantified in replicate experiments, and representative examples are shown in a combined panel (Fig. 1C and 3B).

In liquid YEA, cells released from quiescence were imaged after 24 hours. For Tet-induced Ppn1 complementation, *ppn1*Δ cells treated with AhT to induce Ppn1 expression (as described in $G_0$

induction, survival, and perturbation assays) either 4 hours before release from quiescence or from the time of release were fixed with 70% ethanol, stained with DAPI (Santa Cruz Biotechnology sc-3598) and Calcofluor White (Sigma 18909), and examined on a Zeiss microscope with a 40x objective. DAPI staining revealed chromosome segregation defects, while Calcofluor highlighted aberrant septation and cytokinesis failures. Representative fields are shown in Fig. S3C and S11B, including wild-type and *ppn1*Δ cells during proliferative growth (controls).

***CDK activity analysis by FRET***

CDK activity dynamics during quiescence exit were monitored using the Eevee-spCDK biosensor (*43*). Wild-type and *ppn1*Δ strains carrying the biosensor were arrested for 3 days in EMM–N to establish quiescence. For imaging, 200 µL of cells (~1 OD; ~1 × $10^7$) were added to lectin-coated chambered cover glass (Lab-Tek II, cat. no. 155379, 1.5 mm thickness), allowed to adhere for 30–40 min, washed three times with EMM+N, and overlaid with 500 µL EMM+N. Cells were imaged immediately at 30 °C on an Olympus FV3000 using a 60× oil-immersion objective. Images were acquired every 5 min for 12 h (wild type) or 15 h (*ppn1*Δ). CFP was excited at 445 nm, and emission was collected at 460–500 nm (CFP) and 530–630 nm (YFP), with identical illumination and filter settings applied to all samples.

Images were analyzed in Fiji (ImageJ) as described by (*43*). Background was subtracted by the rolling-ball method (radius = 50 pixels), and single-cell tracking was performed using the LIM Tracker plugin (*61*). Regions of interest (ROIs) corresponded to nuclei identified in CFP images. FRET efficiency was quantified as the YFP/CFP intensity ratio, plotted over time for individual cells. CDK activation onset, peak activity, and deactivation timing were calculated from these traces as in (*43*).

**Aneuploidy (Ch16) assay**

The large linear minichromosome Ch16 (530 kb), carrying centromere 3 and the ade6-M216 allele, has been described previously (*37*); in strains with the host ade6-M210 allele this yields an ade6$^+$ phenotype. For aneuploidy assays, Ch16-containing WT and *ppn1*Δ strains were grown at 30 °C in EMM–Ade, starved of nitrogen to enter quiescence, and maintained for up to three weeks. Weekly aliquots were plated on YE low-adenine medium, where colony color reports Ch16 status. White colonies indicated retention (3:1), whereas red colonies reflected aneuploidy arising during the first mitotic division after quiescence (loss of Ch16 with gain of an endogenous chromosome, >3:0). Half-sectored colonies (missegregation of Ch16 at the first division) and fully sectored colonies (loss in later divisions) were rare but scored separately. For proliferating controls, single colonies of WT or *ppn1*Δ strains carrying Ch16 were grown at 30 °C in YEA and plated on YE low-adenine medium, with overall instability quantified as the combined frequency of red, half-sectored, and sectored colonies.

**$G_0$ suppressor screen**

We performed the genetic screen as described previously (*20*), with minor modifications. Five freshly isolated *ppn1*Δ strains were alternated between proliferation (YEA, 1 day) and quiescence (EMM-N, 3 days) at 30 °C for 25–30 cycles by replica plating. Single colonies were isolated on YEA and assayed for $G_0$ viability to identify suppressors. Genomic DNA from 20 candidates (derived from independent populations) was purified (NucleoSpin Tissue Kit; Takara

740952), pooled into four groups (five per pool), sheared to ~350 bp (Covaris S220), and prepared with the TruSeq DNA PCR-Free LT kit (Illumina). Barcoded libraries were sequenced on a NovaSeq 6000 (Illumina; 150 bp paired-end).

FASTQ files were trimmed with Cutadapt v4.0 (*62*) and aligned to the *S. pombe* reference genome (PomBase; https://www.pombase.org) using Bowtie2 v2.4.5 (*63*). Alignments were processed with SAMtools v1.15.1 (*64*) for conversion, sorting, and indexing.Variants were called with GATK HaplotypeCaller v4.2.6.1 (*65*) and manually filtered to retain only high-confidence, non-synonymous SNPs unique to suppressor pools (expected allele frequency ≥ 20%, parental *ppn1*Δ at 100%). Genes with recurrent mutations or pathways represented by multiple independent hits were prioritized, yielding eight unique suppressor mutations in transcription elongation factors (nine events in total).All candidates were validated by Sanger sequencing, and linkage was tested by two-step backcrossing: SNPs were first isolated in a wild-type background and then reintroduced into a fresh *ppn1*Δ strain. Additional indels, including frameshift alleles in the same or related genes, were detected but not pursued further.

**Chromatin immunoprecipitation (ChIP), ChIP-seq and ChIP-qPCR**
ChIP experiments were performed as described previously (*66*), with minor modifications. Cells were induced into quiescence as above, and 25 OD equivalents were used per ChIP. For RNA Pol II, cohesin, and condensin ChIPs, cells were crosslinked in 3% formaldehyde for 30 min at room temperature with slow rocking and quenched with 2.5 M glycine. For Ppn1 ChIPs, an additional crosslinking step with dimethyl adipimidate (Sigma 285625) was performed for 45 min at room temperature.

Cell pellets were resuspended in 150 µL ChIP lysis buffer (50 mM HEPES-KOH pH 7.5, 140 mM NaCl, 1% Triton X-100, 1 mM EDTA, 0.1% sodium deoxycholate, 1 mM PMSF), disrupted with glass beads using a bead-beater, and sonicated. Lysates were adjusted to 1 mL with lysis buffer, clarified by centrifugation (13,000 rpm, 10 min, 4 °C), and precleared with 20 µL control beads (ChromoTek bab-20) for GFP-trap IPs, or Protein A/G agarose and sepharose beads (Invitrogen 101041 and 15920010) for Ppn1, Cut14, and Pol II IPs (1 h, 4 °C, rotation). After centrifugation (1,000 rpm, 1 min), the supernatant was transferred to a fresh tube; 20 µL was reserved as input control. Immunoprecipitation was carried out using antibodies against RNA PolII Ser2P (Abcam ab5095 and Millipore 04-1571-1) or Anti-V5 tag (Bio-Rad MCA1360), with Protein A/G beads for recovery. GFP-tagged proteins were precipitated using GFP-trap agarose beads (ChromoTek gta-20). Immunoprecipitated DNA and input DNA were analyzed by qPCR or processed for Illumina sequencing.

For ChIP-seq, chromatin was sheared using a Covaris S220 sonicator (peak power 75, duty factor 10, 200 cycles/burst; 15 cycles of 2 min treatment with 1 min delay, 4 °C), and libraries were prepared from immunoprecipitated DNA or input DNA using Ultra II DNA Library Prep Kit (NEB) and sequenced on NovaSeq 6000 Illumina platform. For ChIP-qPCR, chromatin was sheared using a probe sonicator (21% amplitude, 45s on / 2 min off, 4 cycles, 4 °C). DNA from IPs and inputs were quantified using iTaq Universal SYBR Green Supermix (Bio-Rad 1725121) on a CFX96 real-time PCR detection system (Bio-Rad). Relative enrichment was calculated by the ΔΔCt method, using *fbp1* as the reference locus. Oligonucleotides used for ChIP-qPCR are listed in Table S1.

**Proximity-dependent biotinylation (TurboID)**
TurboID-based proximity labeling was performed with full-length or truncated Ppn1 fused at the C terminus to TurboID–3×Myc. Cultures (1 L) were induced into quiescence (EMM–N, 30 °C) for 2 days without biotin supplementation. Cells ($OD_{600} \approx 1.0$; total 100 OD for full-length Ppn1 (FL), and 1000 OD each for ΔC-short (S) and ΔC-long (L)) were harvested, resuspended in fresh EMM–N containing 50 µM biotin, and incubated for 24 hours before extraction. In addition to a no-biotin control, an untagged isogenic wild-type strain, treated identically, served as a control for endogenous biotinylation and non-specific capture. Total protein extracts were prepared as described (*67*) with minor modifications. Cell pellets were resuspended in cold lysis buffer (150 mM HEPES-KOH pH 7.5, 150 mM KCl, 1.5 mM $MgCl_2$, 1 mM EGTA, 1% NP-40, 0.4% sodium deoxycholate, 10% glycerol, 1 mM DTT, 1 mM PMSF, 1× PLAAC, and 1× EDTA-free protease inhibitor cocktail) and lysed with cryogenic grinding in a Mixer Mill (Retsch, MM400). Lysates were adjusted to 5 mL with the same buffer, sonicated (Sonics VibraCell VCX300; 3 × 10 s at 20% amplitude), and treated with Benzonase (5000 U; Santa Cruz Biotechnology sc-202391) for 1hours at 4 °C. Protein concentration was normalized by Bradford assay, and the total extract was incubated with Streptavidin resin (100 µl; GenScript L00353) for 3 hours at 4 °C.

Beads were washed three times with 50 mM Tris-HCl pH 7.5 (twice with 2%SDS, once without SDS), three times with RIPA buffer containing 1 mM DTT (50 mM Tris-HCl pH 7.5, 150 mM NaCl, 1.5 mM $MgCl_2$, 1 mM EGTA, 1% NP-40, 0.1% SDS, 1 mM DTT), and five times with 20 mM ammonium bicarbonate. All washes were 5 min at room temperature with agitation. Bead-bound proteins were denatured in 2 M urea, reduced with 1 mM DTT, alkylated with 5 mM iodoacetamide, digested on-bead with trypsin, and recovered for LC–MS/MS analysis.

**RNA extraction**
Total RNA was prepared from cells (total 10 OD) in quiescence (EMM-N) or activated for cell-cycle re-entry (YEA) at 30 °C using MasterPure Yeast RNA purification kit (Lucigen MPY03100) and was treated with DNase I (Lucigen D9905K) according to manufacturer recommendations. Ribosomal RNAs were depleted using the QIAselect FastSelect - rRNA Removal Yeast Kit (Qiagen 334215). Library construction was performed using TruSeq Stranded Total RNA Prep Kit (Illumina) according to the manufacturer's instructions. Paired-end sequencing was performed on the Illumina NovaSeq 6000 platform.

**Data Analysis**
***Mass Spectrometry (LC–MS/MS Data Analysis)***
**TMT-MS:** Raw LC–MS/MS data were analyzed using Proteome Discoverer **(**v2.2.0.388, Thermo Fisher Scientific Inc.) with the *TMT 6-plex quantification* workflow. Peptide spectral matches were identified by the Sequest HT search engine against the *S. pombe* UniProt database (Taxon ID 284812, 2022). Reporter-ion intensities for the six TMT channels (126–131 Da) were quantified using the *Reporter Ions Quantifier* node, and protein abundances were normalized using the default 6-plex parameters. Only proteins with ≥ 2 unique peptides and present in all three biological replicates of either condition were retained. Grouped abundances for sample and control were used to compute relative abundance and fold change; proteins with $\log_2$ fold change > 1 were considered enriched.

**TurboID-MS:** Label-free proximity-labeling (TurboID) data were processed using MaxQuant v2.6.6.0 under default LFQ settings(*68*). Spectra were searched against the *S. pombe* UniProt database (Taxon ID 284812, 2024), and nonspecific interactors were removed using two controls: (i) tagged strains cultured without biotin and (ii) untagged strains grown with biotin. Proteins identified with ≥ 2 razor + unique peptides and nonzero intensities across both replicates were retained for comparison among full-length Ppn1 (FL), ΔC-short (S), and ΔC-long (L).

For FL vs S, analysis focused on proteins detected in FL but absent in S, corresponding to interactions dependent on the Ppn1 C-terminal region. For S vs L, only proteins shared between FL and S but lost in L were analyzed, identifying contacts requiring the conserved IDR segment. LFQ intensities were $\log_2$-transformed, median-normalized, and missing values imputed with a QRILC-like method. Fold changes were calculated from mean $\log_2$ values, and significance assessed by Welch's t-test. Statistical analyses were performed in Python v3.8 (pandas, NumPy, SciPy(*69*)) and visualization in GraphPad Prism.

***ChIP-seq Data Visualization and Analysis***

Raw paired-end reads were assessed with FastQC v0.11.9, summarized in MultiQC v1.12, and trimmed with Trim Galore v0.6.7 (paired-end, Phred < 20) (*70-72*). Filtered reads were aligned to the *Schizosaccharomyces pombe* reference genome (ASM294v3, GCF_000002945.2) using BWA-MEM v0.7.17, and spike-in reads were aligned to the *Saccharomyces cerevisiae* genome (R64, GCF_000146045.2) (*73*). Alignments were processed with SAMtools v1.15 for conversion, sorting, and duplicate handling (sort n, fixmate -m, coordinate sort, markdup) to preserve fragment-level accuracy (*64*). Spike-in normalization of Psc3–GFP and RNA Pol II (Ser2P) ChIP datasets employed constant *S. cerevisiae* chromatin (from a BY4741 strain carrying a GFP-tagged condensin subunit Brn1) added prior to immunoprecipitation. Uniquely mapped spike-in reads were counted (samtools view -c -F 4), and sample-specific scaling factors were computed as $1 \times 10^6$ / spike-in reads. Genome-wide coverage tracks were generated with deepTools v3.5.1 (bamCoverage, 10-bp bins) using spike-in–derived scaling for quantitative comparison across samples (*74*). In parallel, RPM-normalized tracks were produced by counting properly mapped reads (-F 260), computing coverage with BEDTools v2.30.0 (genomecov -bg), scaling to $1 \times 10^6$ reads, and converting to bigWig with UCSC utilities (*75*).

Unless noted otherwise, peaks for visualization and downstream analyses (including the heatmaps in Fig. 2) were called with MACS2 v2.2.7.1 ($q < 0.05$) and filtered against PomBase-derived blacklists to remove rDNA, centromeric, telomeric, and mitochondrial regions; only peaks with > 2-fold enrichment were retained (*76*). Spike-in–normalized relative enrichment ratios (rER) were computed as ChIP/Input using bigwigCompare (10-bp bins), with low-coverage regions (< 20% of mean input) masked and rER smoothed with a 100-bp centered moving average. Background-subtracted tracks (e.g., NLS-GFP controls) were obtained with bigwigCompare --operation subtract. Genes were ranked by RNA Pol II occupancy in wild-type cells (top 10% = highly expressed; bottom 50% = poorly expressed). Metagene profiles were generated with deepTools computeMatrix (scale-regions; 1 kb upstream, 3 kb gene body, 1.5 kb downstream; 10-bp bins) and visualized with plotProfile. ChIP-seq coverage tracks and peak regions were visualized using Integrative Genomics Viewer (IGV) v2.15 for qualitative inspection and locus-level analysis (*77*).

For the Ppn1 complementation assays, background-subtracted bigWig files were converted to bedGraph with UCSC bigWigToBedGraph, and Psc3–GFP peaks were called with MACS3 v3.0.3 (bdgpeakcall) using a signal-intensity cutoff of 3 (-c 3). Peak–gene assignment considered protein-coding genes from the *S. pombe* GFF3 (PomBase; version matched to alignment), prioritizing overlaps with annotated gene bodies; otherwise, the nearest TSS or TES (transcription start or end site) was assigned based on midpoint-to- boundary distance. To reduce redundancy, the highest-scoring peak per gene per sample was retained. Replicates were processed independently, and only genes assigned in both replicates for a given condition were included in downstream overlap analyses (Venn comparisons).

For spatial concordance analysis, the 127 peaks identified in the full-length Ppn1 (FL) complementation were designated as the reference set. Test datasets (WT, 4h induction, and IDR-only) were compared to this reference using the 'bedtools closest' function to identify the single nearest neighbor. Signed distances relative to the FL peak were converted to absolute base-pair distances, linking each reference peak to at most one peak from each test dataset. Spatial restoration was quantified by classifying FL peaks as matched or unmatched across a series of genomic distance thresholds (0, 50, 100, 250, 500, 1,000, 2,000, and 5,000 bp), with exact overlaps defined as 0 bp (Fig. S9A). Distance variability across individual FL loci (indexed 1–127) was visualized using Python, plotting proximity to 4h and IDR peaks in a mirrored configuration or with a broken y-axis to facilitate direct locus-by-locus comparison of peak proximity (Fig. S9B). Comprehensive gene lists for all conditions (original and newly gained targets) are provided in Table S3.

***RNA-seq Data Analysis***
Adapter-trimmed reads were aligned to the *Schizosaccharomyces pombe* genome (Ensembl release 61; ASM294v2) using STAR v2.7.11b (--quantMode GeneCounts (*78*)) to obtain gene-level counts. Count matrices were processed in DESeq2 (R v4.3.1 (*79*)): Ensembl IDs were mapped to gene IDs with BioMart annotations (*80*), low-count genes were removed prior to normalization, and only annotated protein-coding genes were retained. Differential expression was evaluated by pairwise contrasts in DESeq2, with genes considered significant at adjusted $p < 0.05$. For temporal clustering, we utilized a curated list of 121 canonical G1-S upregulated genes in cycling *S. pombe* as defined previously (*81*). Statistical analyses and data visualization were performed in R v4.3.1 using pheatmap, factoextra, DESeq2, FactoMineR, RColorBrewer, ggpubr, and grid. Gene Ontology enrichment analysis was performed using ShinyGO v0.77 (*82*).

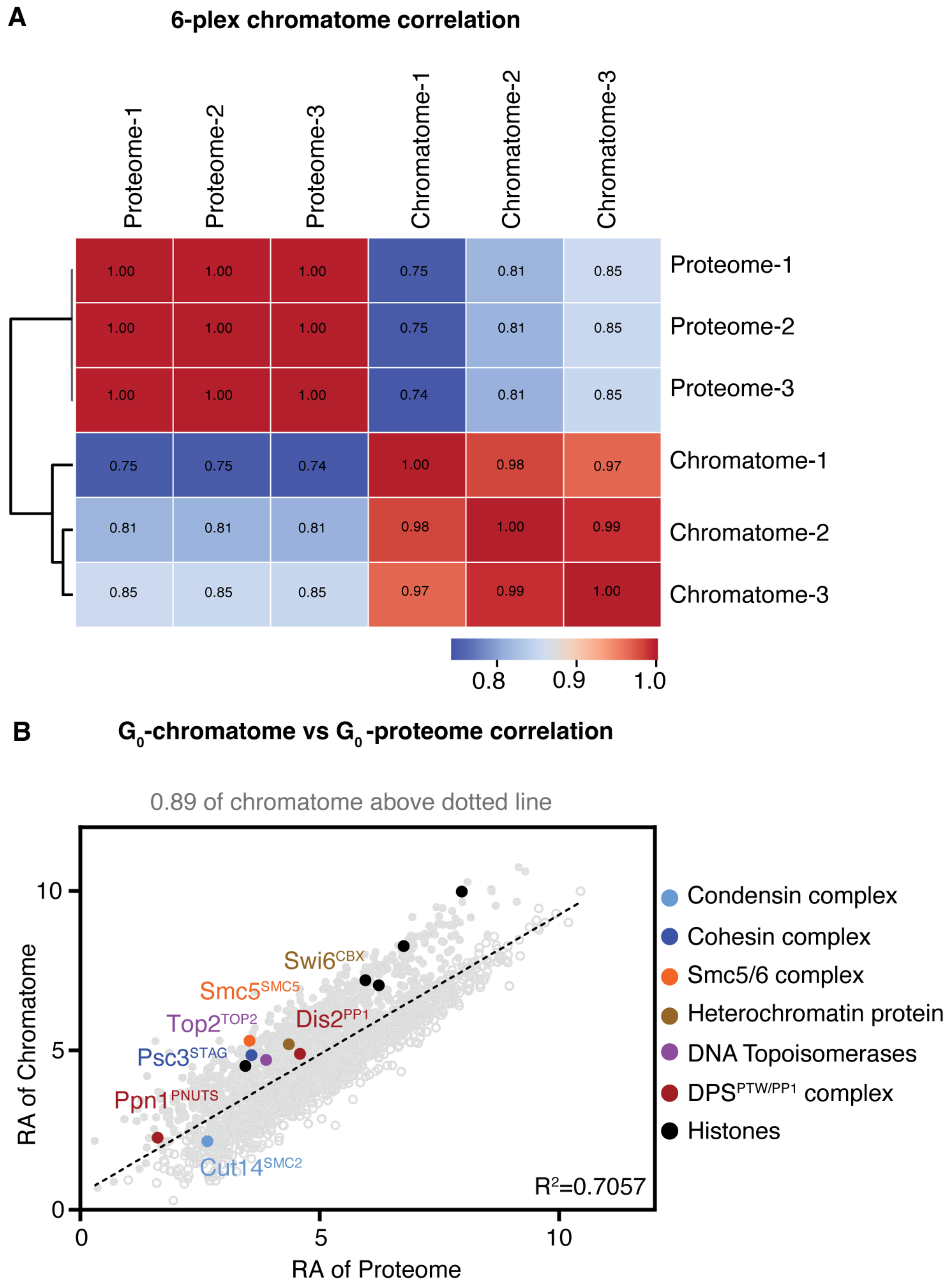


**Fig. S1. The quiescent chromatome is enriched for conserved structural organizers. (A)** Pearson correlation matrix of protein abundances (TMT 6-plex proteomics) comparing whole-cell and chromatin-bound ("chromatome") fractions from quiescent WT cells ($n$ = 3 biological replicates). **(B)** Relative abundance (RA) correlation showing selective enrichment of histones and conserved chromatin organizers within the chromatome. See Table S2 for protein lists.

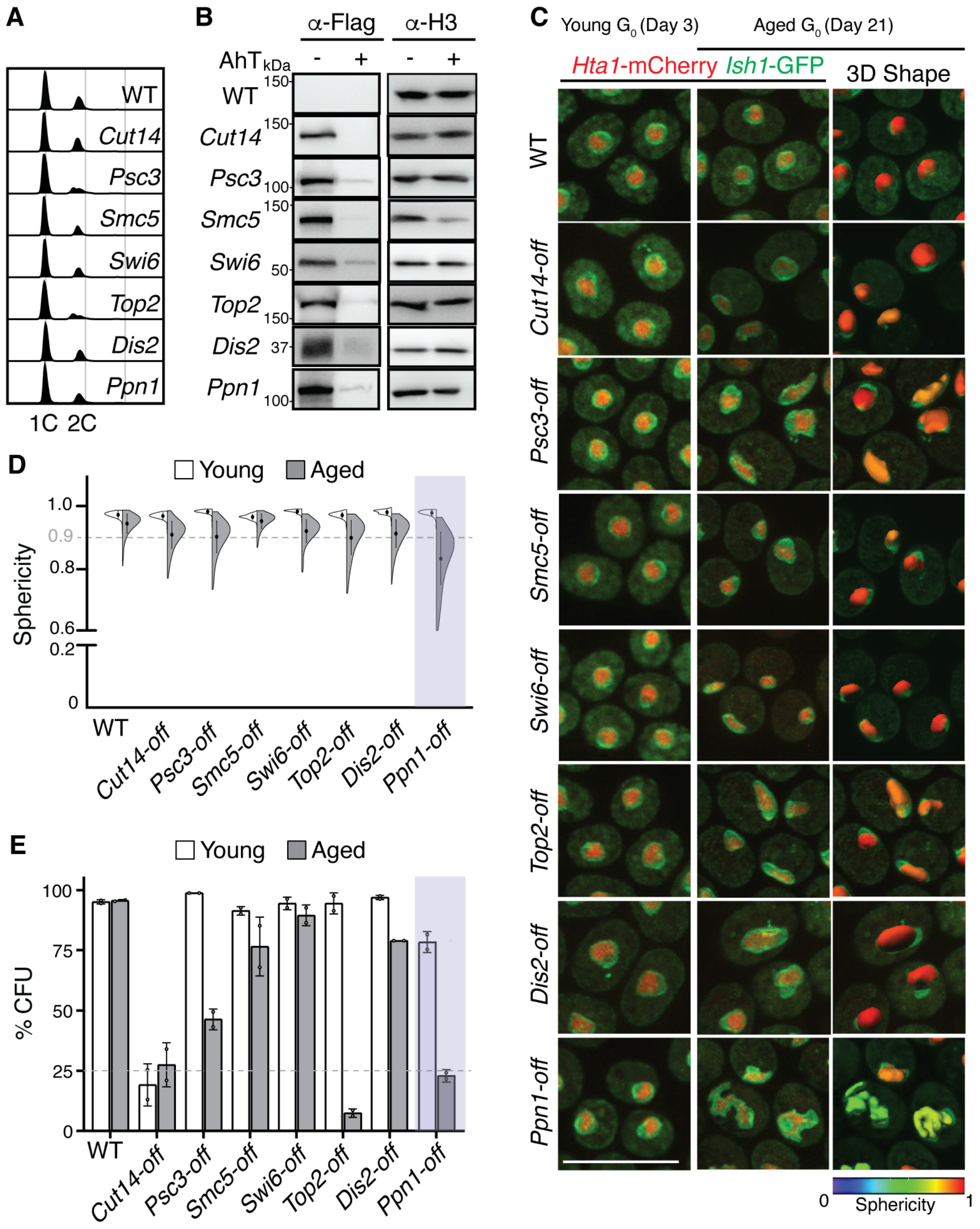


**Fig. S2. Targeted screen identifies Ppn1 as a primary safeguard of $G_0$ chromatin. (A)** Flow cytometry profiles 24 hours post-nitrogen withdrawal confirming normal $G_0$ entry for all Tet-off (AhT-inducible) strains. **(B)** Immunoblot of AhT-dependent depletion of 3xFLAG-tagged candidates after 72 hours in quiescence; Histone H3, loading control. **(C)** Hta1–mCherry (chromatin) and Ish1–GFP (nuclear envelope) imaging in young (Day 3) and aged (Day 21) $G_0$ cells with AhT-dependent depletion. **(D)** Chromatin sphericity quantification (3D shape analysis, $n > 400$ cells, 2 biological replicates) revealing loss of compaction in specific depletion strains with age. **(E)** $G_0$ viability assays identifying Ppn1 as a major quiescence dependency. Data represent 2 biological replicates; CFU, colony forming units; error bars, SD; scale bars, 10 µm.

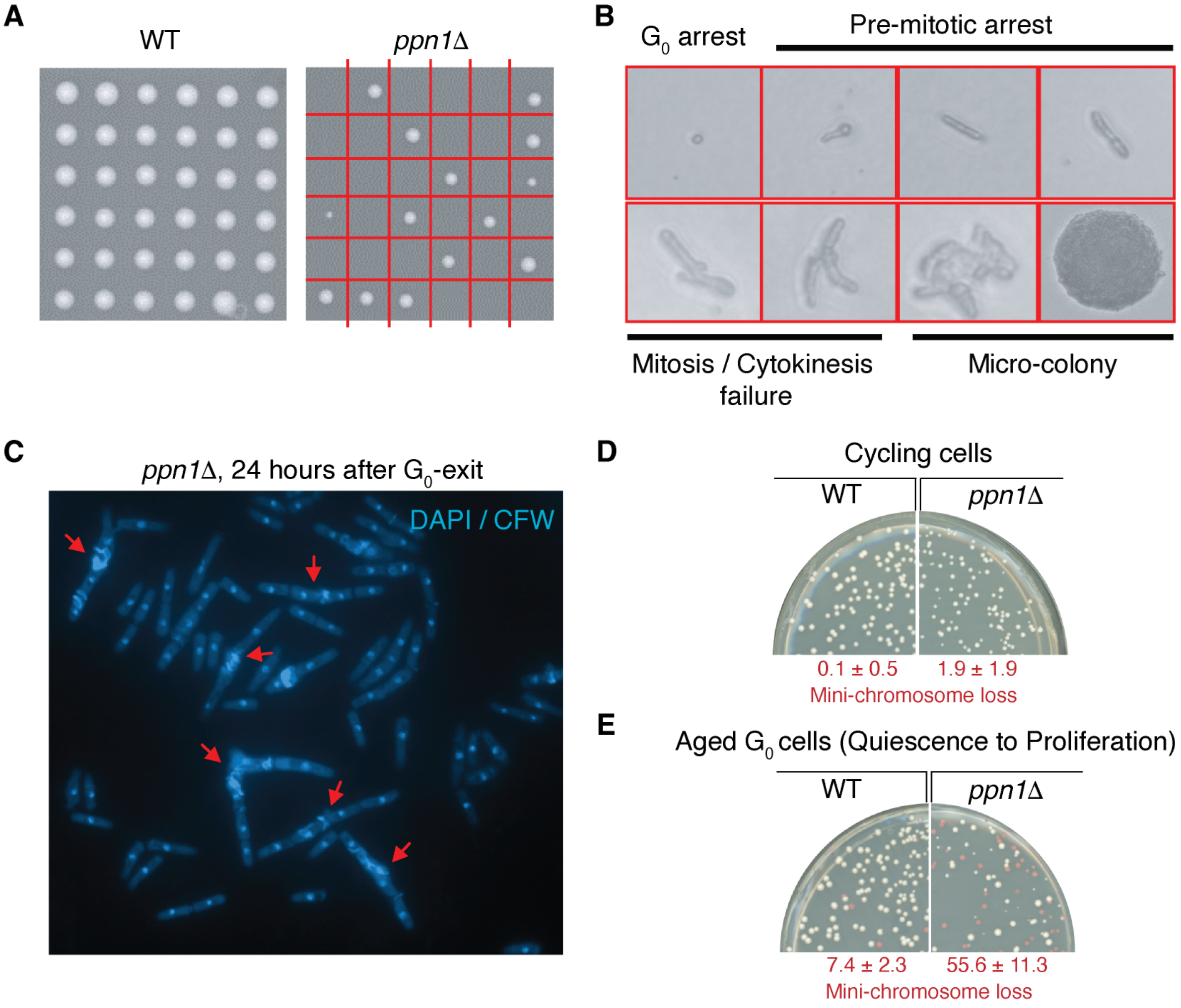


**Fig. S3. Ppn1 loss leads to catastrophic mitotic failure and aneuploidy upon reactivation.** **(A)** Single-cell colony formation tracking of WT and *ppn1*Δ $G_0$ cells. **(B)** *ppn1Δ* terminal morphologies categorized as $G_0$-arrested, pre-mitotic arrested, mitosis/cytokinesis failure, and non-expanding microcolonies. **(C)** DAPI/Calcofluor White (CFW) imaging 24 hours post-release (3-day $G_0$); red arrows highlight chromosome segregation and division defects. **(D)** Minichromosome (MC) assay frequency of red aneuploid colonies among survivors, indicating high-frequency aneuploidy upon quiescence exit ($n \geq 3$ biological replicates).

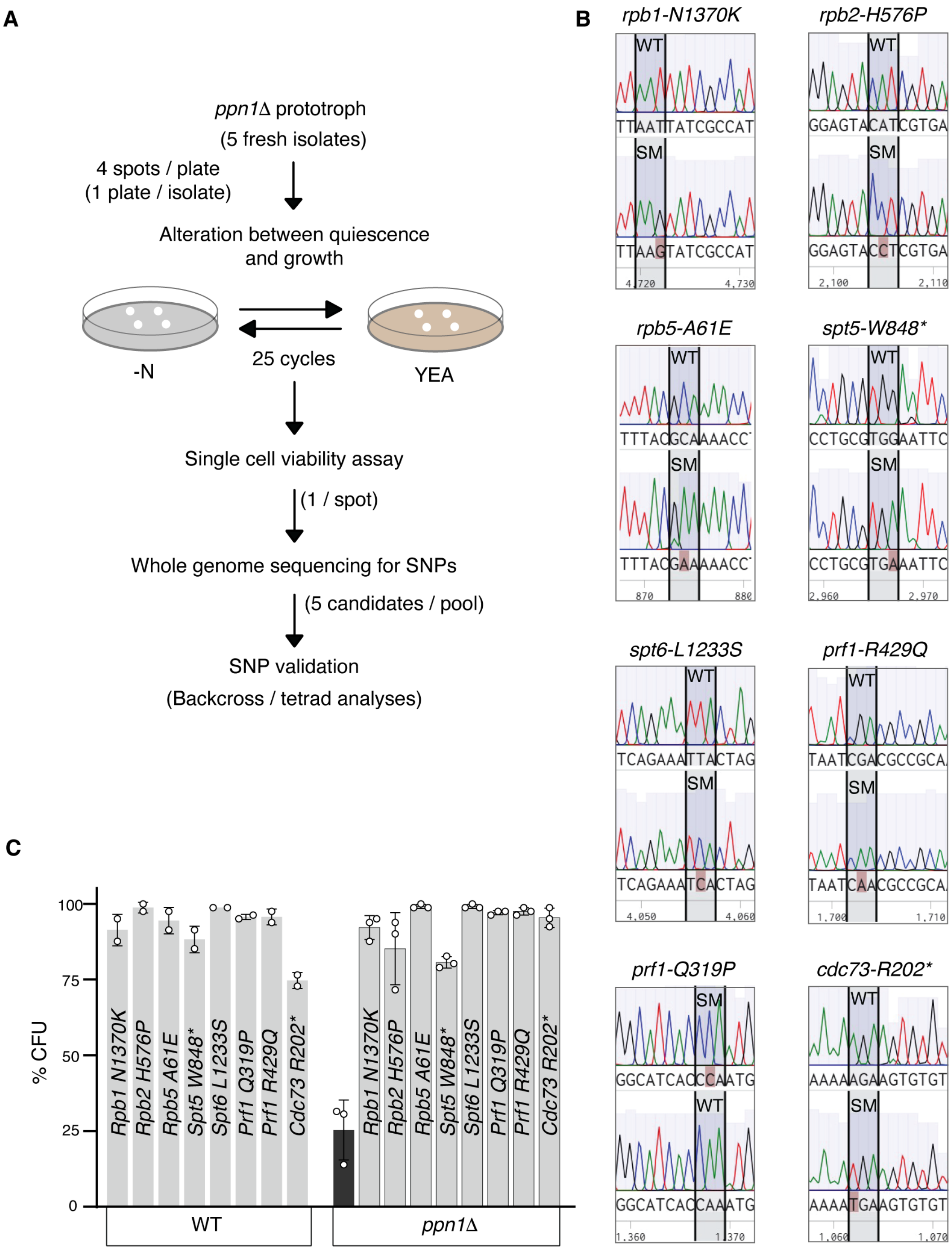


**Fig. S4. Suppression of ppn1Δ $G_0$-defects by mutations in RNA polymerase II and transcription elongation factors. (A)** Suppressor screen workflow; recurrent non-synonymous mutations mapped to essential RNA Polymerase II subunits and conserved positive regulators of transcription elongation. **(B)** Sanger chromatograms of suppressor alleles (SM). **(C)** *ppn1Δ* viability rescue after 14 days in quiescence by individual suppressor mutations. $n \geq 2$ biological replicates; error bars, SD.

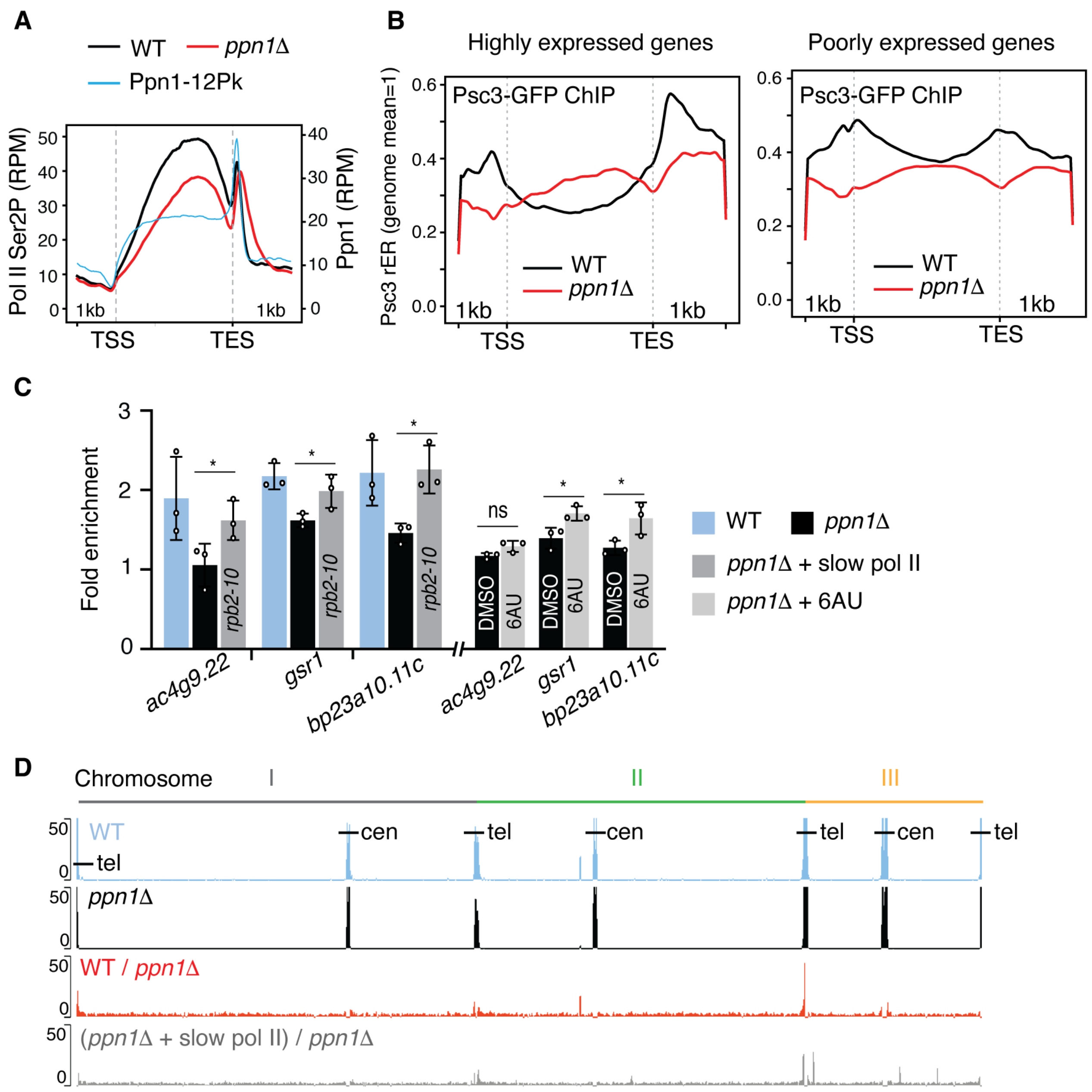


**Fig. S5. Ppn1 prevents transcriptional readthrough to preserve cohesin across the $G_0$ genome. (A)** Metagene ChIP-seq of Ppn1–12PK and elongating Pol II (Ser2P; spike-in normalized; $n$ = 339, top 10 % expressed genes; 3-day $G_0$); Ppn1is enriched at gene ends to prevent downstream Pol II accumulation (TSS, transcription start site; TES, transcription end site). **(B)** Metagene ChIP-seq (3-day $G_0$) of Psc3–GFP showing cohesin loss in *ppn1*Δ across both high and low expression loci. **(C)** ChIP–qPCR confirmation (3-day $G_0$) that dampening elongation (slow Pol II or 6AU-treatment) restores cohesin occupancy at representative 3′ ends (*$P \leq 0.05$; $n$ = 3 biological replicates; error bars, SD). **(D)** Whole-genome views of centromeres and telomeres (*cen*, centromere; *tel*, telomere; I, II and III indicate *S. pombe* chromosomes).

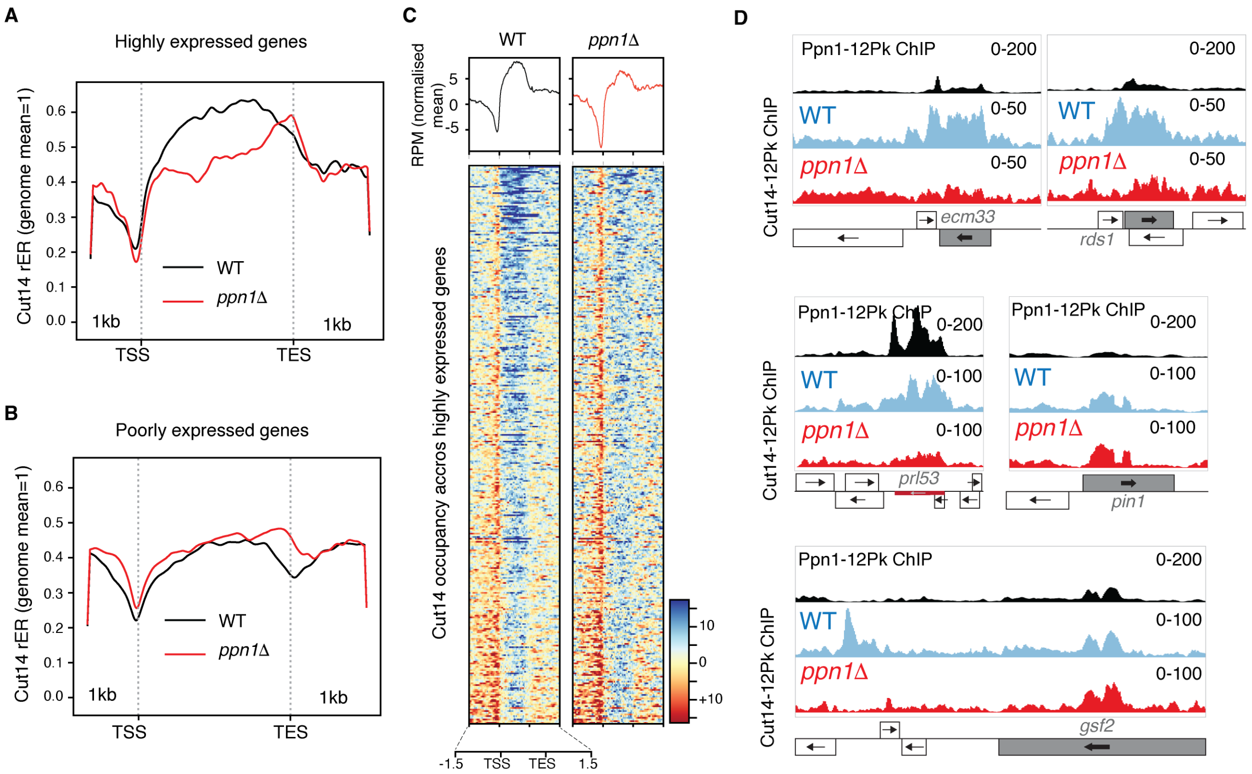


**Fig. S6. Selective condensin loss in ppn1Δ reflect secondary effects, not a direct result of transcriptional readthrough. (A, B)** Metagene ChIP-seq (3-day $G_0$) of condensin (Cut14–12Pk) showing gene-body enrichment upstream of the TES across all expression levels. **(C)** Heatmaps (3-day $G_0$) revealing reduced condensin at multiple gene loci in *ppn1Δ* compared to WT. **(D)** Browser views showing condensin reduction at canonical mitotic targets (*ecm33*, *rds1* and *prl53*) but not at new locations in $G_0$ (*pin1* and *gsf2*).

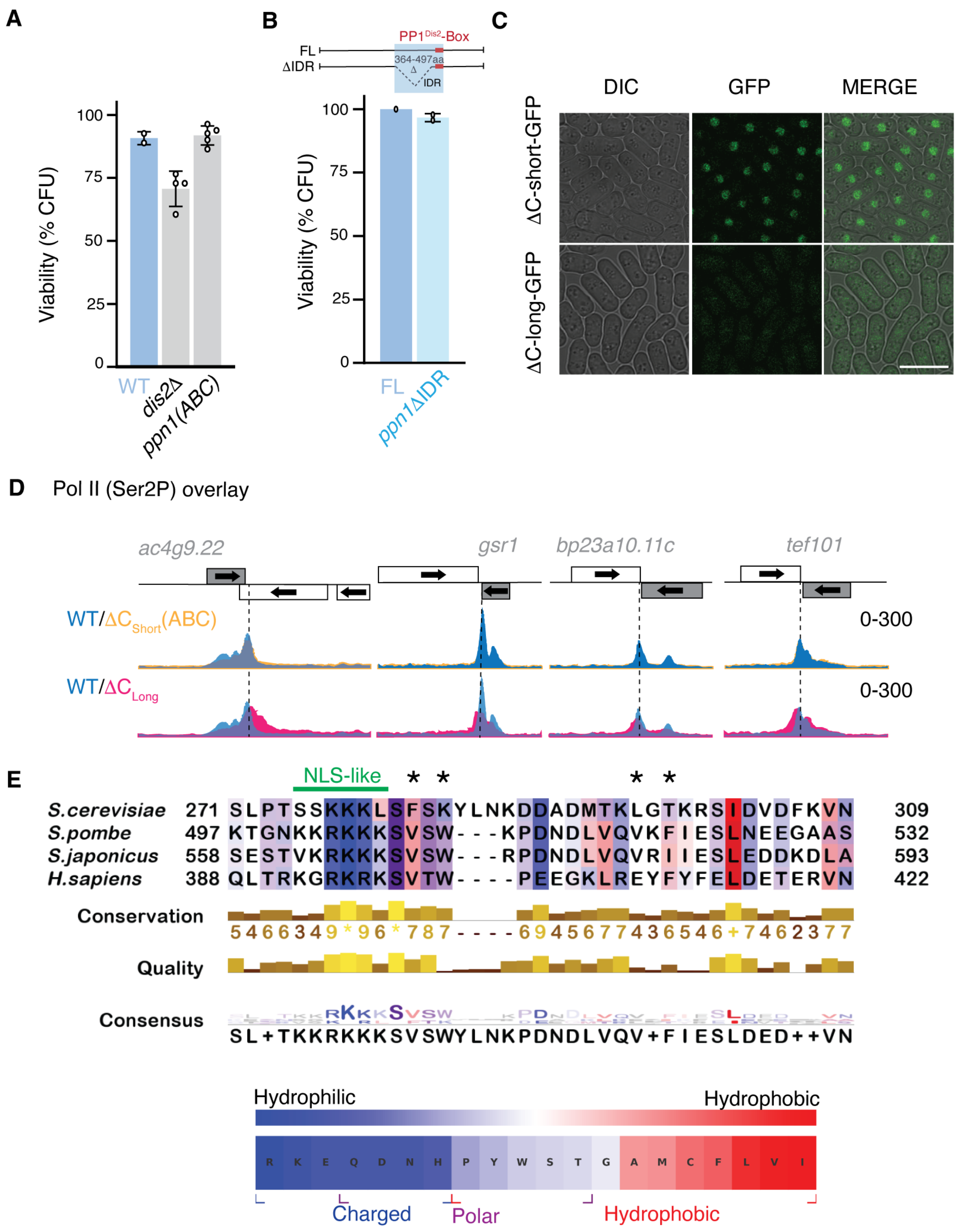


**Fig. S7. A conserved Ppn1 disordered segment drives nuclear function independently of the PP1 phosphatase. (A)** Deletion of the PP1 phosphatase *dis2* or mutation of PP1-binding residues in Ppn1-ABC (V508A, W510A, V519A) does not affect viability in aged $G_0$ cells (21

days), showing that Ppn1's quiescence function is PP1-independent. **(B)** Removal of Ppn1 IDR residues 364–497 does not affect $G_0$ viability over 21 days; only residues 497-532 are essential as shown in Fig. 3B. **(C)** in cycling cells, nuclear localization of the ΔC-short (aa 1–532) variant vs. cytoplasmic retention of ΔC-long (aa 1–496, lacking the critical IDR). **(D)** Ser2P Pol II ChIP-seq (3-day $G_0$) showing pervasive readthrough in ΔC-long, mirroring *ppn1Δ* patterns. **(E)** IDR sequence alignment across eukaryotes highlighting conserved hydrophobic patterning and an NLS-like motif. Asterisks indicate PP1-binding sites; scale bars, 10 μm.

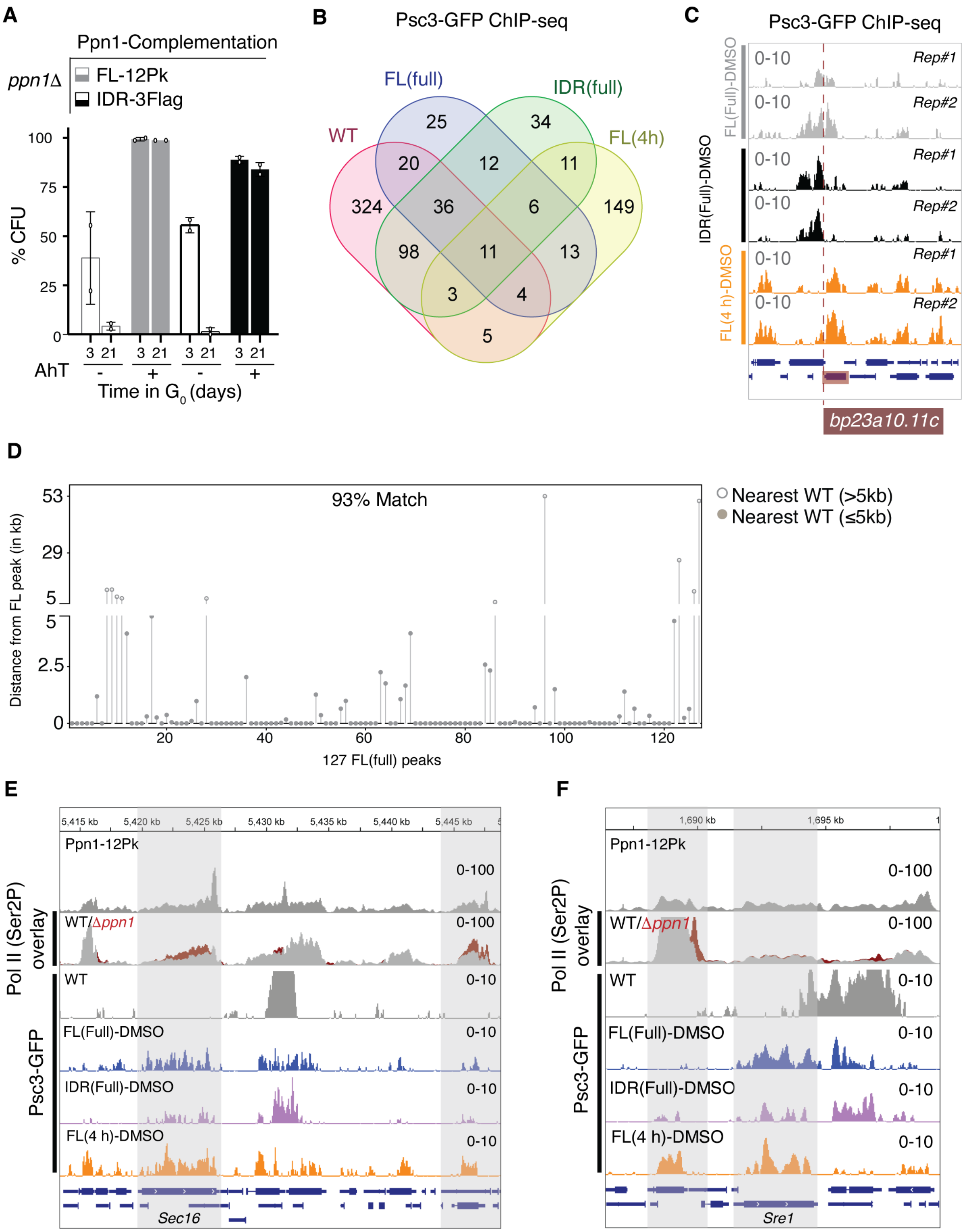


**Fig. S8. The Ppn1 IDR alone restores cohesin stability and viability in quiescent cells. (A)** $G_0$ viability rescue by ectopic Ppn1 IDR (aa 342–532) expression. **(B)** Venn diagrams summarizing the overlap of cohesin-occupied loci between WT (established at $G_0$-entry) and complementation conditions (formed *de novo* in quiescence). See Table S3 for gene lists and binning criteria. **(C)** Browser view of restored Psc3–GFP enrichment at a native target locus in 3-day $G_0$ following continuous Ppn1 full-length (FL) or IDR only induction, or with an acute (4 h) pulse administered prior to harvest (related to Fig. 4); profiles are spike-in–normalized and background-subtracted. Red shading, native WT $G_0$ cohesin targets. **(D)** Spatial mapping of Psc3–GFP peaks in FL (full) relative to WT at 3-day $G_0$. 93% of the 127 reference peaks reside within 5 kb of native WT sites, defining a core architectural template for $G_0$ viability. **(E, F)** Browser views of *de novo* cohesin recruitment (3-day $G_0$). Ppn1–12Pk and Ser2-P Pol II tracks identify active $G_0$ transcription sites. Psc3–GFP tracks show *de novo* enrichment, visualized via spike-in normalization and subtraction of background *ppn1*Δ (DMSO) signal (also related to Fig. 4). Gray shading: active genes with cohesin gain.

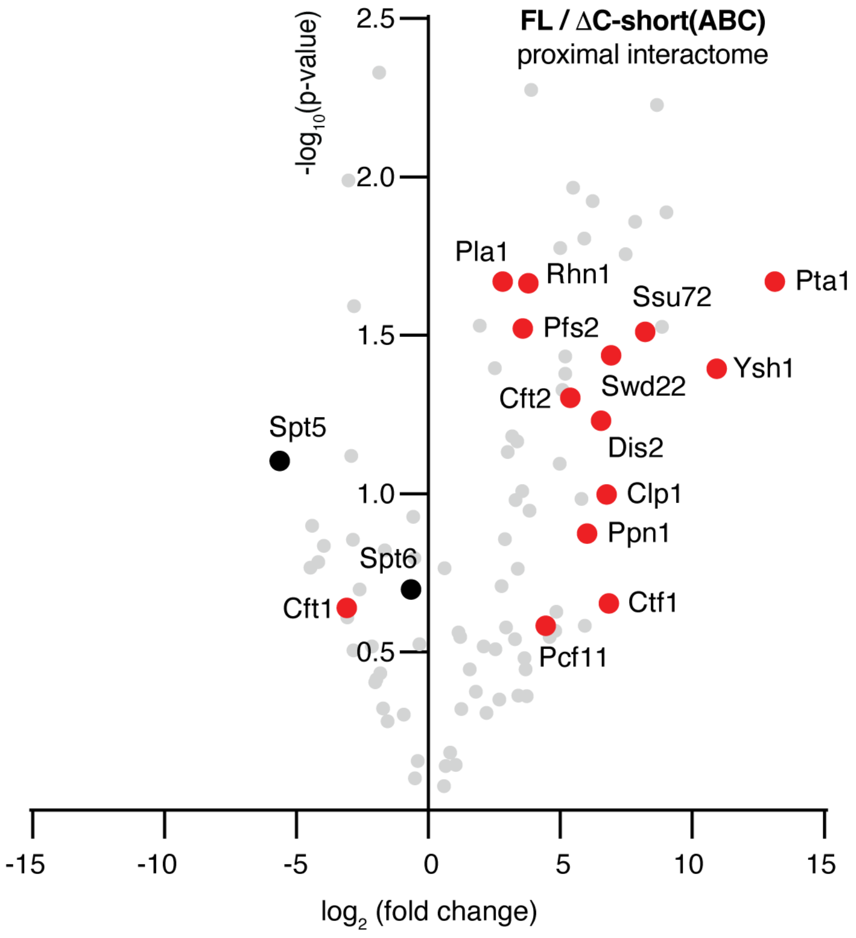


**Fig. S9. Functional ΔC-short(ABC) loses Dis2$^{PP1}$/Swd2.2 (DPS) binding but retains core CPF and Pol II interactions.** Volcano plot of TurboID proximity labeling (3-day $G_0$) comparing functional FLAG-tagged ΔC-short(ABC) with full-length Ppn1 (FL). Persistence of CPF (red) and Pol II (black) contacts explains the variant's ability to support transcription termination in $G_0$ ($n = 2$ biological replicates). Background interactors were removed using no-biotin and untagged controls; only proteins detected with ≥ 2 unique peptides were included. See Table S4 for protein list.

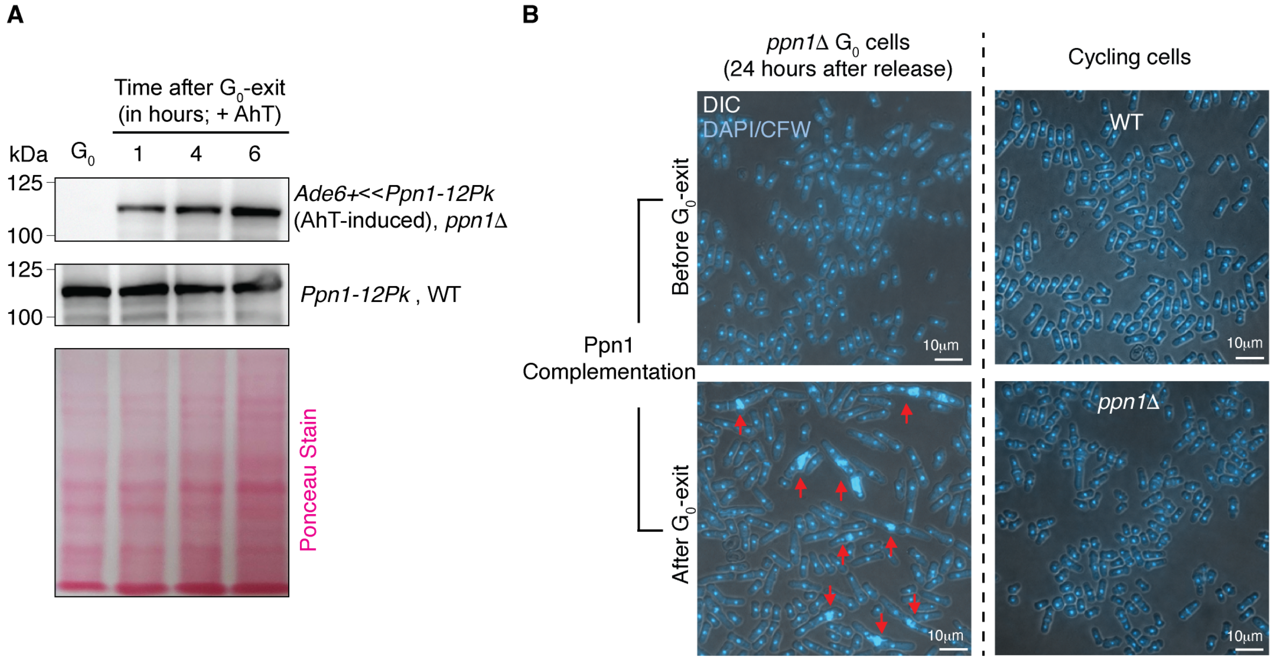


**Fig. S10. Ppn1 restoration in aged $G_0$ cells prevents mitotic errors. (A)** Immunoblot of Ppn1 induction kinetics (3-day $G_0$). *ppn1*Δ cells released into EMM+N show rapid Pk-tagged Ppn1 accumulation post-AhT induction (0–6 hours). WT levels and Ponceau loading controls are shown for reference. **(B)** DAPI/Calcofluor (CFW) imaging of reactivation mitosis, 24 hours post-release (from 3-day $G_0$). Ppn1 induction within $G_0$ (pre-exit) rescues nuclear division, while post-exit induction fails to prevent catastrophic segregation and cytokinesis defects.

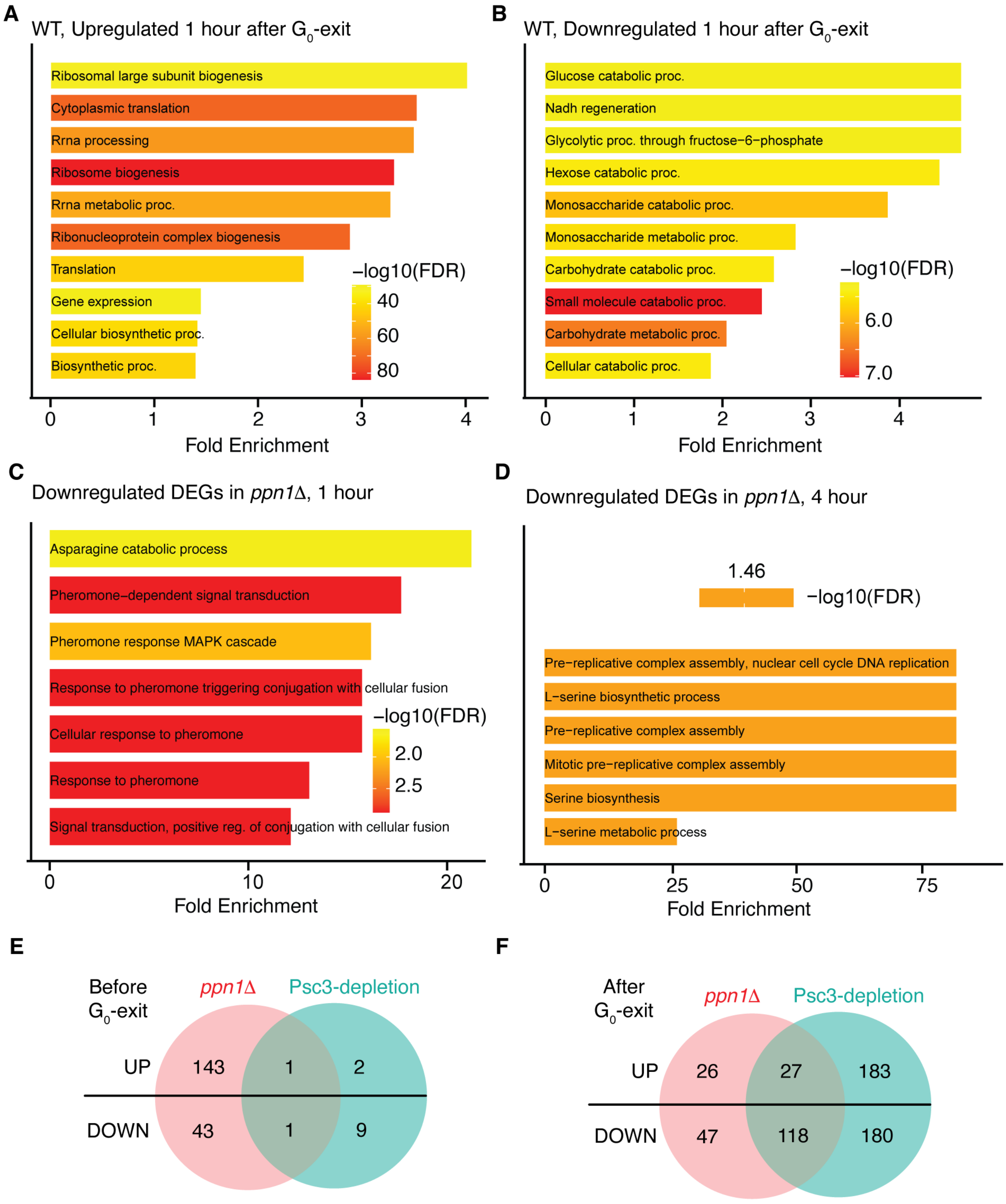


**Fig. S11. Transcriptome dynamics reveal early regulatory disruption in *ppn1*Δ cells. (A, B)** Gene Ontology (GO) analysis from WT transcriptomic shifts one-hour post-release from quiescence (3-day $G_0$; $|\log_2$ fold-change$| \geq 1$, adjusted $P < 0.05$; n = 1971). Early reprogramming features upregulation of biosynthetic and translational pathways and repression of glycolysis, signaling a rapid metabolic transition to growth. **(C, D)** GO analysis of downregulated differentially expressed genes (DEGs) in *ppn1*Δ (Day3 $G_0$). At one-hour post-reactivation, MAPK signaling, centromere assembly, and replication-licensing transcripts are reduced; by 4 hours after $G_0$-exit, downregulation of pre-replicative complex components predominates, consistent with delayed S-phase entry ($|\log_2$ fold-change$| \geq 0.6$, adjusted $P < 0.05$). **(E, F)** Convergence of DEGs in *ppn1*Δ and cohesin-depleted (*Psc3*-off) transcriptomes. Minimal overlap in $G_0$ (<1%) transitions to 66% convergence 4 hours post-reactivation, identifying a shared transcriptional dependency on cohesin integrity ($|\log_2$ fold-change$| \geq 0.6$, adjusted $P <$ 0.05). GO via ShinyGO v0.8; see Table S5 for fold-changes.

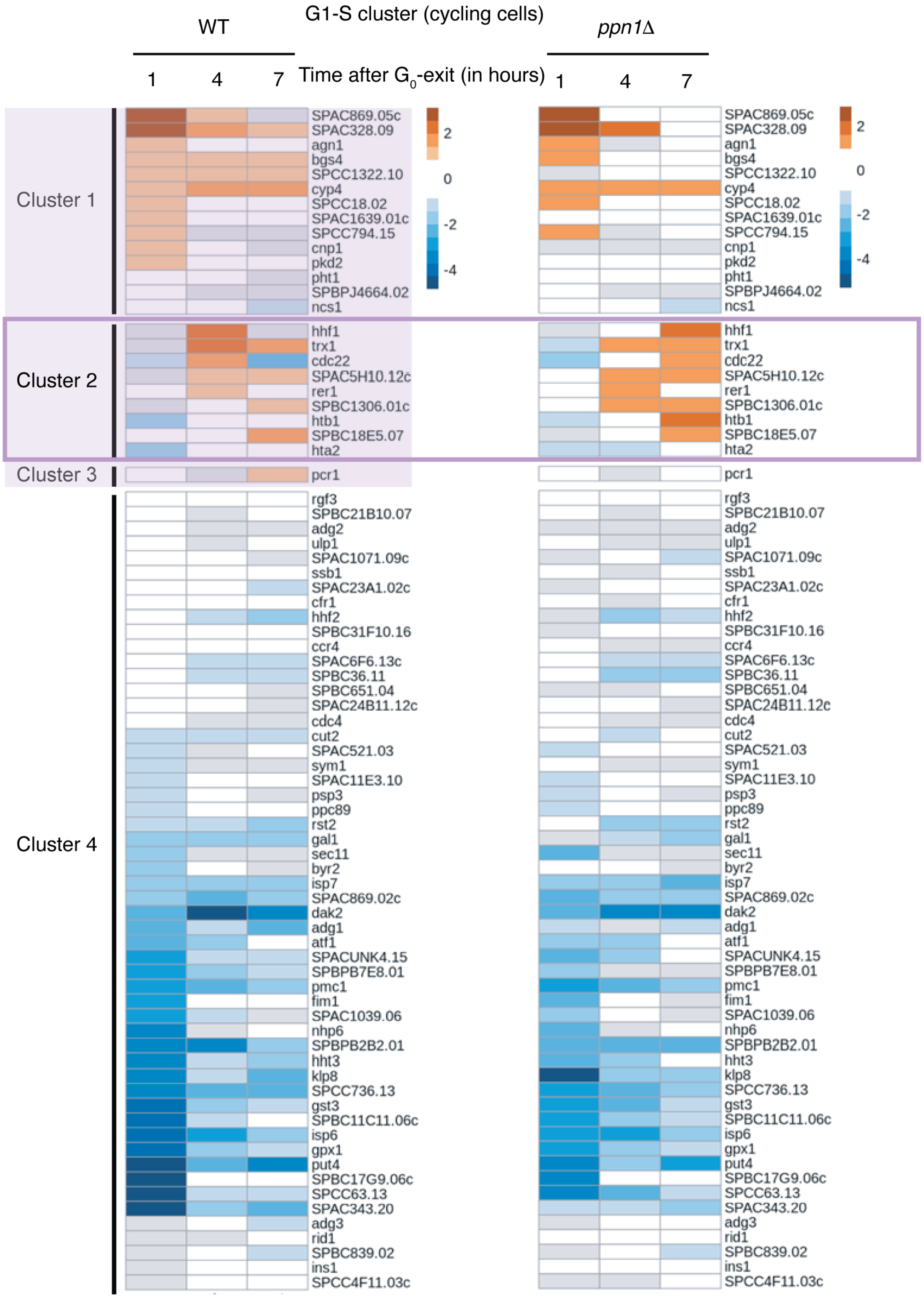


**Fig. S12. Distinct $G_1$–S transcriptional trajectories during quiescence exit in 3-day $G_0$.** Temporal clustering of 121 canonical genes defined as $G_1$–S-upregulated in cycling *S. pombe*

(see Materials and Methods, *RNA-seq Data Analysis*). While these genes drive the cycling $G_1$–S transition, ~80% remain stable in both WT and *ppn1*Δ, suggesting that $G_0$ exit is transcriptionally distinct from post-mitotic $G_1$–S activation. However, within the subset of genes normally activated during exit, *ppn1*Δ exhibits significant misexpression and kinetic delays, exemplified by a 3-hour peak shift in Cluster 2 histone genes (from 4 hours in WT to 7 hours in *ppn1*Δ).

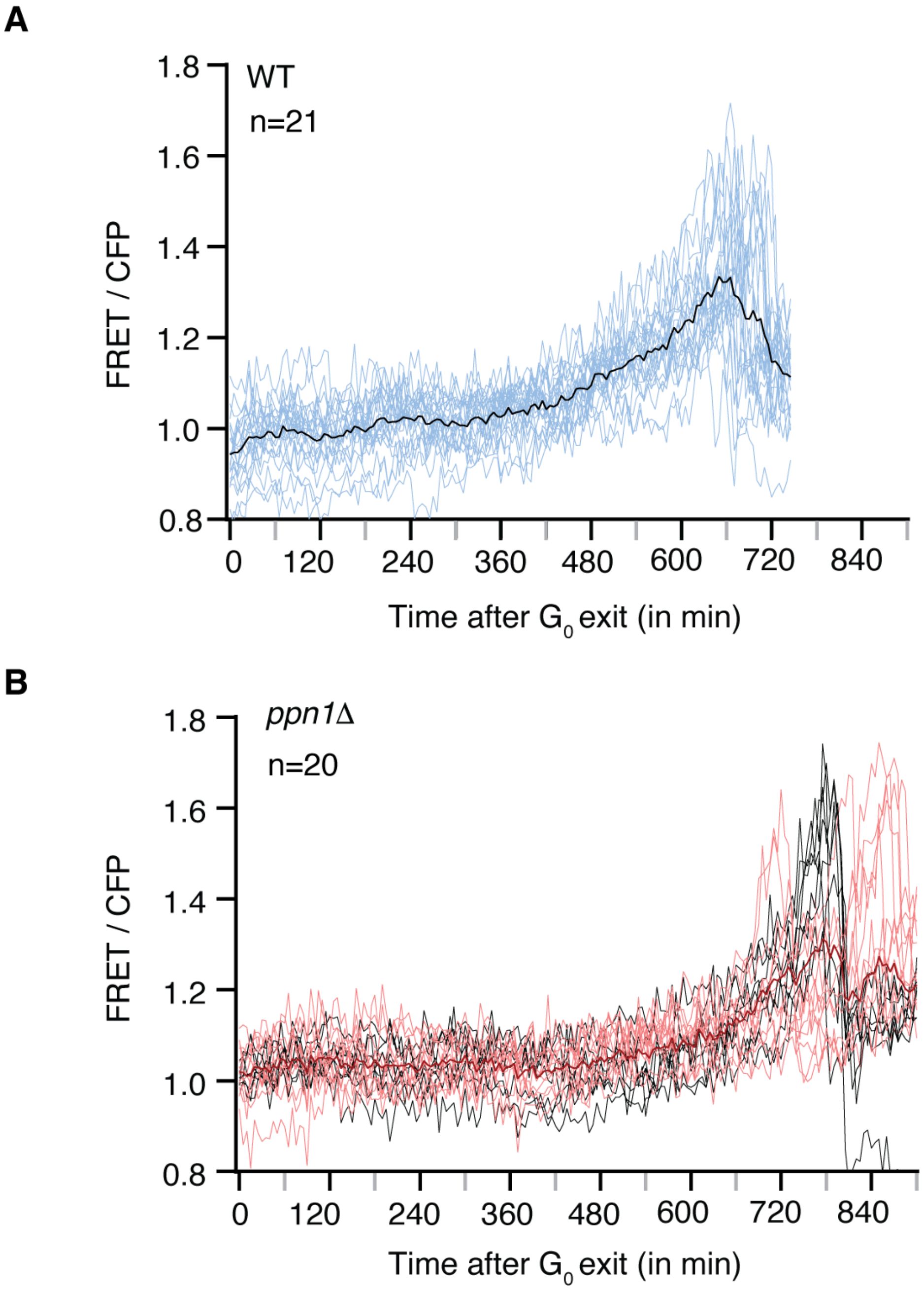


**Fig. S13. CDK activity dynamics during re-entry are distorted in ppn1Δ cells.** Single-cell CDK activity traces (Eevee-spCDK FRET biosensor) during re-entry from 3-day $G_0$. **(A)** WT cells display coordinated CDK activation through the first mitosis. **(B)** *ppn1*Δ traces reveal pronounced heterogeneity (black and red) and aberrant oscillations around mitosis, consistent with impaired CDK coordination and reduced division competence. Each line represents a single-cell trajectory.

**Table S1 | Strains, oligonucleotides, and plasmids used in this study.**

**Table S2 | TMT-based proteome and chromatome datasets in quiescence.** Related to fig. S1.

Normalized TMT reporter intensities from whole-cell proteome (P) and chromatin-bound (C) fractions of quiescent WT cells (3-day $G_0$). Sheet 1 ("TMT 6-plex intensities") provides raw quantitative metrics, including unique peptide counts, protein coverage (%), and peptide-spectrum matches (PSMs). Sheet 2 ("$G_0$-enriched (C > P)") identifies the 3-day $G_0$ chromatome defined by a |log_2FC > 1| (C vs. P) enrichment, including matched human homologs.

**Table S3 | Cohesin occupancy landscape at $G_0$ entry and during architectural rescue.** Related to Fig. 4, C and D, and fig. S8, B and D.

Summary of Psc3–GFP ChIP-seq peaks identified via MACS3 and assigned to the nearest gene summit. WT ($G_0$ entry) is scored by presence/absence. Complementation datasets—FL (full), FL (4 h), and IDR (full)—report *ppn1*Δ-subtracted signal intensity binned by enrichment: ++ (>100), + (<100), or – (absent). The table includes both native WT sites and *de novo* sites gained during Ppn1 full-length/IDR restoration.

**Table S4 | Ppn1 proximity interactome defined by TurboID.** Related to Fig. 3F and fig. S9.

Filtered interactors for full-length Ppn1 and variants ΔC-short (functional) and ΔC-long (non-functional). Data represent two biological replicates filtered against negative controls (untagged + biotin; tagged - biotin). Only proteins with ≥ 2 razor + unique peptides and nonzero intensities in both replicates are included. Metrics provided: unique peptide counts, protein coverage (%), and relative enrichment used for volcano plot generation in Fig. 3F and fig. S9.

**Table S5 | Differential gene expression analysis and heatmap matrices.** Related to Fig. 5, A and B, and fig. S11.

Transcriptomic contrasts across four worksheets. Sheets 1–2 report DESeq2 outputs (mean expression, |log_2FC|, and adjusted P-value) for *ppn1*Δ and *Psc3*-off cells in $G_0$. Sheets 3–4 provide input matrices for reactivation heatmaps, listing differentially expressed genes in WT at one-hour and 4 hours post-exit (|log_2FC| > 1, Padj < 0.05) with cross-referenced |log_2FC| values for *ppn1*Δ and *Psc3*-off; non-significant shifts are denoted NA.
